# Defining the role of RNase E in the mycobacterial degradosome-like network

**DOI:** 10.64898/2025.12.24.696391

**Authors:** Abigail R. Rapiejko, Ying Zhou, Vidhyadhar Nandana, Junpei Xiao, Jared M. Schrader, Scarlet S. Shell

**Affiliations:** Department of Biology & Biotechnology, Worcester Polytechnic Institute, Worcester, Massachusetts, 01609, USA; Department of Biological Sciences, Wayne State University, Detroit, Michigan, 48202, USA. Current affiliation: Nuclear Dynamics and Cancer Program, Cancer Epigenetics Institute, Fox Chase Cancer Center, Philadelphia, Pennsylvania, 19111, USA; Program in Bioinformatics & Computational Biology, Worcester Polytechnic Institute, Worcester, Massachusetts, 01609, USA; Department of Biology, Indiana University, Bloomington, Indiana, 47405, USA

**Keywords:** RNase E, mRNA degradation, mRNA decay, mRNA processing, degradosome, degradosome-like network

## Abstract

mRNA degradation is a fundamentally important process that is regulated in response to stress in the globally important pathogen *Mycobacterium tuberculosis*. Several mycobacterial ribonucleases (RNases) are hypothesized to function together to coordinate mRNA degradation, but the interactions among them are mostly undefined. One of the rate-limiting enzymes, RNase E, contains intrinsically disordered regions (IDRs). Here, we have defined the interactions among major mycobacterial mRNA degradation enzymes and identified the functions of the two IDRs of RNase E in the nonpathogenic model *Mycolicibacterium smegmatis*. We found that the two IDRs differentially impact mRNA degradation rates *in vivo* but are largely functionally redundant in their impact on steady-state transcript abundance. *In vitro*, the IDRs are uninvolved in catalysis but play major roles in RNA binding and interactions with other mRNA degradation enzymes, namely PNPase, RNase J, and RhlE1, including stimulation of RhlE1’s activity. *In vivo*, these enzymes localize with RNase E, but its IDRs play only a minor role, suggesting substantial redundancy in subcellular localization mechanisms. Collectively, we propose a degradosome-like network model in mycobacteria, held together by dynamic, transient interactions among RNA degradation enzymes and RNA that can be disrupted during physiologically relevant stress to allow for adaptability.

**GRAPHICAL ABSTRACT:** 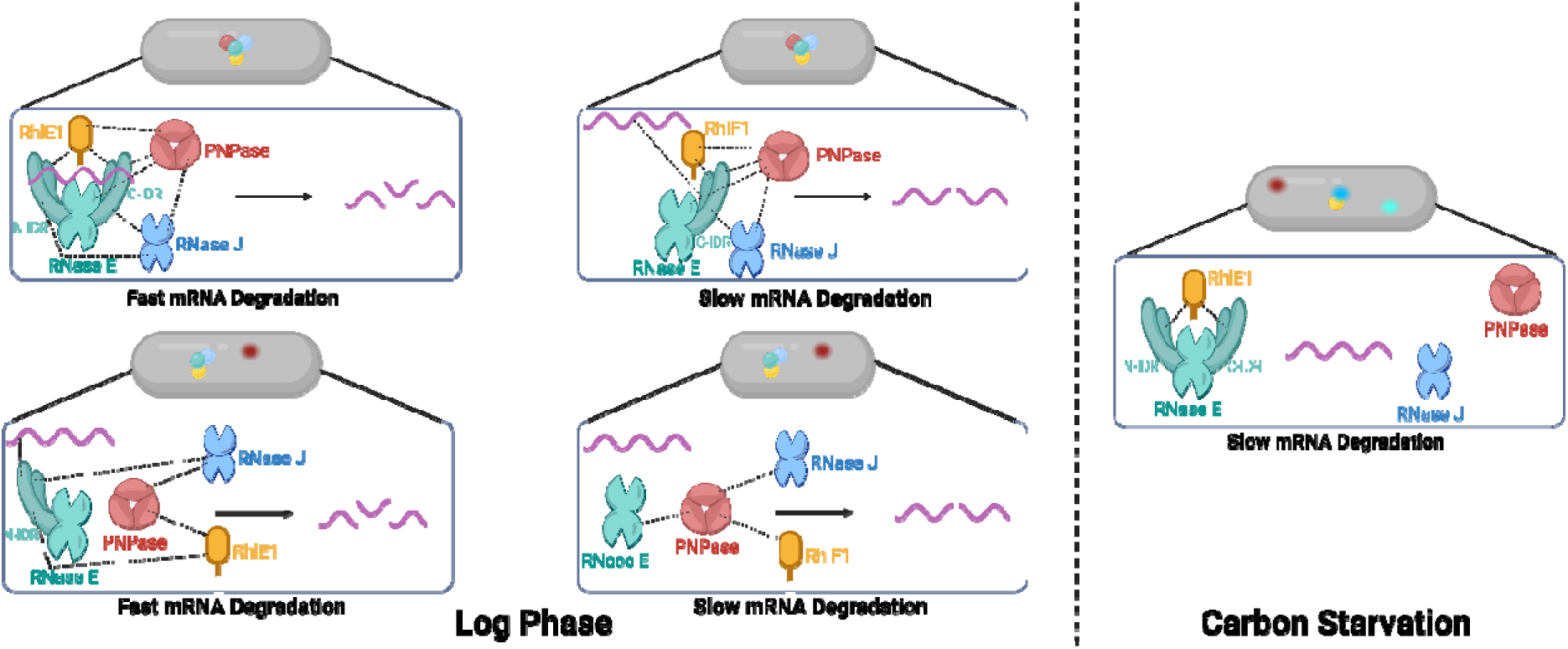

## INTRODUCTION

Mycobacteria are a clinically relevant group of bacteria that includes *Mycobacterium tuberculosis,* the causative agent of tuberculosis, which causes millions of reported cases and deaths each year (1). During *M. tuberculosis* infections, the bacteria face several stressors from the human host such as hypoxia, low pH, and reactive oxygen and nitrogen species (reviewed in (2)), yet the bacteria can still survive and often tolerate antibiotics. One way that mycobacteria respond to energy stressors like hypoxia is global stabilization of their transcriptomes by decreasing mRNA degradation rates (3,4), as it is energetically expensive to synthesize and degrade RNA. However, this stabilization cannot be explained by changes in the abundance of RNA processing and degradation enzymes (3,5). The mechanisms by which mycobacteria post-translationally regulate their RNA processing and degradation enzymes are unknown.

To understand the regulation of mRNA degradation, it is necessary to define the components and organization of the molecular machinery that carries out this process. Endoribonucleases, exoribonucleases, and RNA helicases are the principal enzymes that carry out mRNA degradation. In mycobacteria, the endoribonuclease RNase E, exonuclease PNPase, and dual endo/exoribonuclease RNase J have established roles in this process (6–9).

The essential endoribonuclease RNase E plays a rate-limiting step in the degradation of at least 80% of mRNAs in the nonpathogenic model organism *Mycolicibacterium smegmatis* (10), and its essentiality and large transcriptional impact in *M. tuberculosis* suggest that it has a similarly import role (11). Structurally, RNase E is known to bind magnesium for efficient catalysis of RNA (6) and produces a 5’ monophosphate and 3’ hydroxyl upon RNA cleavage (12). In *Escherichia coli,* zinc binding is required for multimerization of RNase E (12,13) to form dimers, which are needed to form the catalytic interface, as well as tetramers (dimers of dimers) (6). RNase E cleaves long but not short RNAs efficiently when multimerized as tetramers or larger species suggesting a post-translational regulatory mechanism (14).

In Proteobacteria, RNase E has a long C-terminal intrinsically disordered region (IDR) that mediates interactions with other proteins (15,16). However, the IDR of RNase E in *Caulobacter crescentus* is required for an additional noncatalytic function: formation of liquid-liquid phase-separated biomolecular condensates (17). Condensates represent a mechanism of cytoplasmic organization by the formation of membraneless organelles in both bacteria and eukaryotes (reviewed in (18,19)). In *C. crescentus*, RNase E condensates allow for rapid assembly and disassembly of RNA degradation hubs for efficient degradation when needed and adaptability during stress (17,20–22). The condensates formed by RNase E can also recruit other enzymes to promote their activity in *C. crescentus* (20). Strikingly, mycobacterial RNase E has two long IDRs flanking a central catalytic domain (15), suggesting potential additional functionalities in condensate formation and/or specific protein-protein interactions compared to the better-studied Proteobacterial RNase Es which have single IDRs.

Other relevant RNA processing and degradation enzymes in mycobacteria include RNase J, PNPase, and RhlE. RNase J is a dimer with dual endo- and 5’ to 3’ exoribonuclease activities (7) and has gained attention because its deletion in *M. tuberculosis* leads to multidrug tolerance (23,24). It has a non-essential role in rRNA maturation and a specialized role in degradation of a subset of highly structured mRNAs (7,24). PNPase is trimer with 3’ to 5’ exonuclease activity (8,9) that can also act as a polymerase in the reverse direction to add 3’ poly-A tails, which in bacteria generally facilitate mRNA degradation (reviewed in (25)). It is essential in *M. tuberculosis* and *M. smegmatis* (11). Notably, mycobacteria lack two 3’ to 5’ exonucleases (RNase R and RNase II) that have redundant roles with PNPase in *E. coli*, where PNPase is not essential (26,27). RhlE is a DEAD Box RNA helicase that is ATP-dependent and proposed to participate in mRNA degradation in mycobacteria (11).

In some bacteria, multiprotein complexes of RNA degradation proteins termed RNA degradosomes have been described. These are mediated by specific interactions and thus are distinct from condensates of RNA degradation proteins, which are formed by many weak, low-specificity interactions. However, it is likely that degradosomes and condensates can coexist, with proteins such as RNase E participating in both types of interactions simultaneously (22). Degradosomes are thought to be sites of accelerated RNA degradation due to the close proximity of related enzymes as well as sites of allosteric activation mediated by direct protein-protein interactions in some cases (16,28,29). In the classical model of *E. coli* mRNA degradation, cleavage by RNase E is the initiation step, while the helicase can unwind secondary structure so that the exonucleases can finish the job by processive cleavage (30,31). The core *E. coli* degradosome has been well-defined, with RNase E serving as a scaffold that binds RhlB (an RNA helicase), PNPase, and enolase with defined stoichiometries (31–36). These stoichiometries can be impacted by factors such as the presence of RNA and other degradation proteins associated with the core degradosome components at sub-stoichiometric levels that may vary depending upon condition (36). For example, the composition of the degradosome can change in response to stress such as cold shock (37). RNase E binds its partner proteins via a long C-terminal IDR (15,16,38), and is membrane-bound in *E. coli* (39).

Variations of the degradosome have been described in other bacteria including *C. crescentus*, *Bacillus subtilis*, and *Helicobacter pylori*. In *C. crescentus,* RNase E binds aconitase and PNPase using its C-terminal IDR and interacts with either of two DEAD-box RNA helicases (40): RhlB during log phase growth and RhlE during cold shock (41). *B. subtilis* lacks RNase E but encodes the endoribonuclease RNase Y, which is considered a functional equivalent. RNase Y facilitates direct and indirect interactions among PNPase, RNase J1, RNase J2, CshA (an RNA helicase), enolase, and phosphofructokinase (42) using its IDR (43). This association of proteins appears less stable than the *E. coli* degradosome; for example, the *B. subtilis* degradosome was first defined using chemical crosslinking (43) and the subcellular localization pattern of RNase Y at the membrane does not match the cytoplasmic distribution of other degradosome components (44). A complex of these components with defined stoichiometry has not been defined. The *B. subtilis* RNA degradation proteins have therefore been described as forming a degradosome-like network involving transient protein-protein interactions (42–46). In *Helicobacter pylori*, an RNase J-based degradosome has been proposed in which it interacts with RhpA (an RNA helicase) and associates with the membrane independently from RNase Y (47).

There is clearly great diversity in the composition and organization of RNA degradosomes across different species. The mycobacterial RNA degradosome has yet to be directly defined. Immunoprecipitation and mass spectrometry experiments have implicated RNase E, PNPase, RNase J, and RhlE as being potential degradosome members in *M. tuberculosis* (11), but the design of these experiments did not distinguish direct interactions from indirect interactions mediated by other proteins or RNA. The only direct interaction tested and confirmed was between PNPase and RNase J (11). Beyond that single interaction, no direct evidence has previously shown the arrangement and composition of a mycobacterial degradosome.

Here, we use the nonpathogenic model organism *M. smegmatis* to investigate the roles of the IDRs of mycobacterial RNase E as well as define the protein-protein interactions that organize the major mycobacterial RNA degradation proteins. We found that the IDRs of RNase E have major and partially redundant roles in RNA binding, which is reflected in the phenotypic impacts of their deletion in cells. *In vitro,* the IDRs of RNase E also have roles in condensate-like body formation, mediate interactions with three other RNA degradation proteins, and stimulate the activity of RhlE1. Remarkably, these functions only have minor impacts on subcellular localization of the RNA degradation machinery *in vivo*. Other components of the RNA degradation machinery also have direct physical interactions, which may underlie the robustness of their localization patterns in cells. Localization of the major mRNA degradation proteins became more diffuse in response to energy stress, consistent with the idea that subcellular localization may contribute to global regulation of mRNA degradation rates. Together, our findings suggest that mycobacterial RNA degradation proteins are organized into a degradosome-like network that is tunable during clinically relevant stress conditions.

## MATERIALS AND METHODS

### Bacterial Strains and Culture Conditions

Middlebrook 7H9 liquid medium supplemented with 5 g/L bovine serum albumin fraction V, 2 g/L glucose, 0.85 g/L NaCl, and 4 mg/L catalase, 0.2% glycerol, 0.05% Tween 80 was used to grow *M. smegmatis* Mc 155 and its derivatives. Middlebrook 7H10 solid medium supplemented with 5 g/L bovine serum albumin fraction V, 2 g/L glucose, 0.85 g/L NaCl, and 4 mg/L catalase, 0.5% glycerol, was used to grow *M. smegmatis* Mc 155 and its derivatives. Liquid cultures were grown at 37°C with 200 rpm shaking. Carbon starvation media for microscopy was composed of 34.2 µM EDTA, 6.8 µM CaCl_2_ x 2 H_2_O, 491.9 µM MgCl_2_ x 6 H_2_O, 0.83 µM Na_2_MoO4 x 2 H_2_O, 1.8 µM CoCl_2_ x 6 H_2_O, 0.80 µM CuSO_4_ x 5 H_2_O, 5.1 µM MnCl_2_ x 4 H_2_O, 7.0 µM ZnSO_4_ x 7 H_2_O, 180 µM FeSO_4_ x 7 H_2_O, 8.9 mM K_2_HPO_4_, 70.8 mM NaH_2_PO_4_, 15 mM (NH_4_)_2_SO_4_, and 0.05% tyloxapol in water. Antibiotic concentrations for *M. smegmatis* were 25 μg/mL kanamycin, 150 μg/mL hygromycin B, 40 μg/mL nourseothricin, and 1 μg/mL apramycin.

All *M. smegmatis* strains used in this study are listed in Supplemental Table 1, and all *M. smegmatis* plasmids are listed in Supplemental Table 2. For experiments shown in Figures 2, 4A-E, 6, and 7, a single copy of the indicated version of the *rne* gene was present at an L5-integrated plasmid expressed from its native promoter and 5’ UTR (436 nt upstream of the native start codon). These were constructed by first integrating into strain Mc^2^155 a copy of *rne* with its native promoter/UTR and an N-terminal 6xhis-3xFLAG tag into the L5 site on a kan-marked plasmid, then deleting all but the last 150 nt of the native *rne* coding sequence by two-step homologous recombination. Finally, the resulting strain was transformed with nourseothricin-marked L5-integrating plasmids encoding the desired RNase E variants (N-terminal his-FLAG tags and N-terminal mCherry fusion with full-length RNase E or various domain deletions), and colonies in which the nourseothricin-marked plasmid replaced the kan-marked plasmid were identified.

For experiments shown in Figure 4G-H, full-length RNase E or the indicated RNase E IDRs were expressed as fusions with mCherry from the native *rne* promoter and 5’ UTR from Giles-integrating plasmids in strains where the native copy of *rne* was intact.

For experiments shown in Figures 6-7, eGFP-tagged PNPase or Dendra2-tagged RhlE1 or RNase J were expressed from the UV15 promoter and 5’ UTR (48) from Giles-integrating plasmids.

Lennox Luria-Bertani broth and solid media were used to grow *E. coli*. Antibiotic concentrations used for *E. coli* were 50 μg/mL kanamycin, 250 μg/mL hygromycin B, 60 μg/mL nourseothricin, and 20 μg/mL chloramphenicol. *E. coli* NEB-5-alpha (New England Biolabs) was used for cloning. HiFi Assembly (New England Biolabs) was used to make all plasmids for this study.

### RNase E Variants Growth Assay

RNase E variants were grown to mid-log phase (OD_600_ of ∼0.5) then diluted to an OD_600_ of 0.05 in 5 mL. Every 3 hours for 9 hours, a 1 mL aliquot was taken from the shaker, with each tube only being removed for approximately 10 seconds. Cultures were read by OD_600_, diluted in duplicate, and 5 μL spots were plated in duplicate (four spots total) on drug-free plates. Colonies were counted on day 2 and additional colonies were counted on day 3. The four spots were averaged, and CFU/mL was normalized to time 0 and reported. Significance testing was done by comparing the slopes of the line for each variant to full-length in GraphPad Prism.

### RNA Half-lives

RNA extraction, cDNA synthesis, and quantitative PCR to determine mRNA half-lives were done as described in (3).

### RNASeq

Strains were grown in triplicate to an OD_600_ of ∼0.42, and RNA was extracted as described in (3). rRNA was depleted, and Illumina-compatible RNAseq libraries were constructed as described in (49,50). Libraries were sequenced at the UMass Medical School Deep Sequencing core facility on an Illumina HiSeq instrument to obtain 100 nt paired end reads. Reads were aligned to the NC_008596 reference genome using Burrows-Wheeler Aligner (51). The FeatureCounts tool was used to assign mapped reads to genomic features. DESeq2 (52) was used to assess changes in gene expression. Genes with an adjusted p-value less than 0.05 and log_2_ fold changes >1 or <-1 were regarded as differentially expressed. Venn Diagrams were made with BioVenn (53). Raw and processed data are available on GEO, accession number GSE313996.

### Microscopy

Nikon CSU W spinning disk confocal microscope with a CFI 60 plan apochromat λ D 100X oil immersion objective was used to obtain microscopy images. 1 mL of exponential phase *M. smegmatis* cultures were washed once and resuspended in less than 200 µL of 7H9 without dextrose, catalase, or BSA. 10 µL of cells were spotted onto 1.2% agarose pads (w/v agarose in water) and imaged in a chamber at 37°C. Brightfield channel used a 300 ms exposure time. mCherry and mScarlet were imaged with a 561 nm excitation laser and a ET605/52m emission filter. mCherry used 80% laser power and 6 s exposure time, and mScarlet used 80% laser power and 3 s exposure time. Dendra2 and eGFP were imaged using a 488 nm excitation laser and a ET525/36m emission filter. For Dendra2, 80% laser power and 200 ms exposure time were used, and for eGFP, 80% laser power and 100 ms exposure time were used. Images were acquired, denoised, and deconvoluted using Nikon Elements software (5.42.06).

For rifampicin treatment, exponential phase cultures were treated for 30 minutes with 100 µg/mL rifampicin at 37°C shaking at 200 rpm.

Image analysis was performed using ImageJ with the FIJI plugin. For cell length measurements, straight lines were drawn through cells, and the measure tool was used. For assessment of colocalization, segmented lines were drawn through the cell in the brightfield channel, and the plot profile was plotted in both fluorescent channels. The intensity in the eGFP or Dendra2 channel was plotted versus the mCherry channel, and the Pearson’s R correlation value was calculated for each cell in GraphPad Prism. Supplemental Figure 11A-D shows examples of intensity versus distance plots and correlation plots for localized and non-localized cells. Supplemental Figure 11E shows the p-value from Pearson’s R correlations from an example set of 150 single- and double-tagged cells, demonstrating more statistically significant correlations in double-tagged than single-tagged strains. For assessment of the number of fluorescence clusters and coefficient of variation, segmented lines were drawn through cells in the brightfield channel, and the plot profile feature was used in the fluorescent channel. Clusters were defined as instances where the signal intensity plotted along the length of the cell rose above and then dipped below the mean intensity (see Supplementary Figure 6A for an example). The number of clusters in each cell was divided by the cell length to calculate clusters/micron. For all microscopy analyses, 50 cells were counted in each biological replicate, and there were three biological replicates per condition.

### CRISPRi Growth Curve to test PNPase Functionality

Exponential phase cultures were diluted to OD 0.005 and treated with either water (negative control) or 50 ng/mL anhydrous tetracycline (ATc) to induce the CRISPRi system in a final volume of 200 µL in a 96 well plate. The plate was read by an EPOCH2 microplate reader (BioTek®) using Gen5 version 3.04 BioTek® software at 37°C with continuous orbital shaking at 1807 cpm 1mm at 600 nM every 10 minutes for 2 days. Biological triplicates were analyzed.

### Rifampicin Killing Curve to test RNase J Functionality

Biological triplicate strains were diluted to an OD_600 nm_ of 0.1 in 5 mL of media with or without a final concentration of 12 µg/mL rifampicin. 5 µL spots were diluted and plated in duplicate on drug-free plates every day for 5 days. Colonies were counted on day 2 and additional colonies were counted on day 3. The average of the spots was reported, and CFU/mL was normalized to time 0 and reported.

### CRISPRi Growth Assay for Lysates

*M. smegmatis* cultures were grown to mid-log phase (OD_600 nm_ of ∼0.5) then diluted to an OD_600 nm_ of 0.05 in 10 mL. 50 ng/mL anhydrous tetracycline to induce the CRISPRi system or water as a negative control was added to each culture. Every 3 hours for 12 hours, a 1 mL aliquot was taken from the shaker, with each tube only being removed for approximately 10 seconds. Cultures were read by OD_600 nm_. Biological triplicates were analyzed.

### Cell Lysate Cleavage Assays

10 mL *M. smegmatis* cultures were grown for 9 hours with from starting OD_600 nm_ of 0.05 with or without 50 ng/mL anhydrous tetracycline (CRISPRi or repressible RNase E) and pelleted by centrifugation for 10 minutes at 3,900 rpm and 4°C. Cells were washed with 10 mL of 7H9 containing no BSA, dextrose, or catalase 3 times then resuspended in 1 mL of 20 mM tris-HCl pH 7.9, 100 mM NaCl, 5% glycerol, 0.01% IGEPAL, 0.1 mM DTT, 10 mM MgCl_2_ and 1X Halt Protease Inhibitor Cocktail, EDTA-Free (ThermoFisher). Cells were lysed with a FastPrep-24 bead-beater (MP Biomedicals) using 4 cycles of 30 sec at 6.5 m/s, 1 minute on ice between cycles in tubes containing 100 µm zirconium beads (OPS Diagnostics, PFMB 100-100-12). Lysates were centrifuged for 10 min 13,000 rpm at 4°C and the supernatants were collected. Pierce BCA Protein Assay Kit (ThermoFisher) was used to determine total protein concentration in the lysate.

For CRISPRi and RNase E repression lysate cleavage assays, a final concentration of 200 nM of 29 nt FAM-labeled RNA purchased from Integrated DNA Technologies (Supplemental Figure 12A and D-F, Supplemental Table 3) and 400 ng of total protein from the lysate was added to cleavage assay reactions with 20 mM tris-HCl pH 7.9, 100 mM NaCl, 5% glycerol, 0.01% IGEPAL, 0.1 mM DTT, and 10 mM MgCl_2_. Reactions were incubated at 37°C in a thermocycler for 1 hour (PNPase and RNase J) or 2 hours (RNase E), stopped using a final concentration of 12.5 mM EDTA pH 8.0 in formamide, and heated at 70°C for 10 minutes. Samples were run on a 20% urea-PAGE gel then imaged on an Azure 600 imager using the Cy3 channel (524 nm excitation). Two (PNPase and RNase J) or three (RNase E) biological replicates of each strain were grown and lysed. Substrate and product band intensities were calculated using ImageJ with the FIJI plugin and used to calculate the percent of substrate remaining. Ordinary one-way ANOVA with Sidak’s multiple comparisons test was used for significance testing.

For hypoxia lysate cleavage assays, a 22 nt FAM-labeled RNA was used (Supplemental Figure 12G, Supplemental Table 3) at a final concentration of 50 nM with cells grown in hypoxia bottles as described in (3) for 3 days prior to harvesting. The CRISPRi protocol described above was modified using 3.8 µg total protein, then the reactions were stopped after the indicated amount of time. Duplicate strains were grown and lysed. Substrate and product band intensities were calculated using ImageJ with the FIJI plugin and used to calculate the percent of substrate remaining. Simple linear regression was used for significance testing.

For carbon starvation lysate cleavage assays, the 22 nt FAM-labeled RNA was used (Supplemental Figure 12I, Supplemental Table 3) at a final concentration of 50 nM with cells that were grown in carbon starvation media for 24 hours prior to harvesting using the protocol described in (3). The hypoxia protocol was modified using 4.8 µg total protein.

### Western Blotting of Cell Lysates

15 mL *M. smegmatis* cultures were grown to exponential phase and pelleted by centrifugation at 10 minutes at 3,900 rpm and 4°C. Cells were washed with an equal volume of 7H9 containing no BSA, dextrose, or catalase three times then resuspended in 300 µL of 20 mM tris-HCl pH 7.9, 100 mM NaCl, 5% glycerol, 0.1 mM DTT, 0.01% IGEPAL, and 1X Halt Protease Inhibitor Cocktail, EDTA-Free (ThermoFisher). Cells were lysed with a FastPrep-24 bead-beater (MP Biomedicals) using 4 cycles of 30 sec at 6.5 m/s, 1 minute on ice between cycles in tubes containing 100 µm zirconium beads (OPS Diagnostics, PFMB 100-100-12). Lysates were centrifuged for 10 min 13,000 rpm at 4°C and the supernatants were collected. Pierce BCA Protein Assay Kit (ThermoFisher) was used to determine total protein concentration in the lysate, and 3 µg of total protein was loaded onto an SDS-PAGE gel then transferred to a PVDF membrane. Membranes were stained with LiCor Revert 700 Total Protein Stain (LiCor, 926-11021) following the manufacturer’s instructions and imaged to verify equal loading. The membrane was blocked with TBS (20 mM tris-HCl pH 7.6, 137 mM NaCl) with 3% nonfat dry milk (w/v) for 30 min at room temperature, washed with TBS three times, and incubated with monoclonal anti-FLAG M2 mouse antibody (1 µg/mL) (Sigma-Aldrich, F1804) in TBS 3% nonfat dry milk (w/v) overnight at 4°C. Membranes were washed with TBS three times then incubated with goat anti-mouse HRP antibody for 45 minutes at room temperature. The membrane was washed 6 times for 5 minutes each with TBS with 0.05% Tween-20 (v/v) then once with TBS (no Tween). Membranes were developed with Azure Radiance ECL (AC2204) in an Azure 600 imager.

### Overexpression and Purification of Recombinant RNase E Variants

RNase E variants were recombinantly expressed and purified using *E. coli* BL21(DE3) pLysS transformed with pET42-derived plasmids (Supplemental Table 4). 2 L cultures were grown at 37°C and were shaken at 200 rpm to an OD_600 nm_ of 0.5-0.8. Cultures were induced with 1 mM IPTG at 28°C and were shaken at 200 rpm for 4 hours prior to harvest by centrifugation. Cell pellets were resuspended in 10 mL of buffer (20 mM tris-HCl, 150 mM NaCl, 5% glycerol, 0.01% IGEPAL, 10 mM imidazole) with 1x Halt Protease Inhibitor Cocktail, EDTA-Free (ThermoFisher), 40 mg of lysozyme, and 16 U Turbo DNase (Invitrogen). Cells were lysed with a BioSpec Tissue-Tearor (6 cycles of speed 6 then 4 cycles of speed 9 that were 30 s each with 30 s on ice between cycles). Lysates were cleared by centrifugation with 8 mL His-Pur nickel–nitrilotriacetic acid resin 50% slurry (ThermoScientific) and the NaCl concentration in the lysate was increased to 1 M before incubation for 60 minutes on at room temperature with end-to-end rotation. The resin was washed three times with 10 mL of 20 mM tris-HCl, 1 M NaCl, 5% glycerol, 0.01% IGEPAL, and 20 mM imidazole then eluted with 4 mL of 20 mM tris-HCl, 150 mM NaCl, 5% glycerol, 0.01% IGEPAL, 150 mM imidazole with 1x Halt Protease Inhibitor Cocktail, EDTA-Free (ThermoFisher) three times, on end-to-end rocker for 10 minutes each. Eluates were concentrated with Microcon PL-100 (full-length) or PL-30 (other variants) (100,000 or 30,000 NMWL) protein concentrators (MilliporeSigma) to a volume of about 300 µL. Samples were loaded onto 1 cm diameter, 38 ml Sephacryl S-400 (full-length, N-deletion, and C-deletion) or S-200 (N- and C-deletion) High Resolution resin (GE Healthcare) size exclusion chromatography columns run with a BioLogic LP chromatography system (BioRad). A flow rate of 0.25 mL/minute was used, and 500 µL fractions were collected using SEC buffer (20 mM tris-HCl, 150 mM NaCl, 5% glycerol, 0.01% IGEPAL, 1 mM DTT, 1 mM EDTA). Fractions were combined, concentrated, and buffer exchanged using the same Microcon protein concentrators with 20 mM tris-HCL, 100 mM NaCl, 5% glycerol, 0.01% IGEPAL, and 0.1 mM DTT. Representative protein preps for RNase E are shown in Supplemental Figure 2A-E.

### Overexpression and Purification of Recombinant Other Degradosome proteins

PNPase, RhlE1, and RNase J variants were expressed and purified as described above for RNase E, scaled down to 1 L growth (Supplemental Table 4 with corresponding *M. smegmatis* gene IDS in Supplemental Table 5). Microcon PL-30 (30,000 NMWL) protein concentrators (MilliporeSigma) were used for all proteins, and Sephacryl S-400 (PNPase) or S-200 (RNase J and RhlE1) High Resolution resin (GE Healthcare) were used for size exclusion chromatography. Representative protein preps are shown in Supplemental Figure 2F-L.

### Cleavage Assays

To test for stimulation of RNase E by PNPase (Supplemental Figure 9B), a hairpin RNA oligo purchased from Integrated DNA Technologies (Supplemental Table 3) at a final concentration of 316 nM was incubated with 100 nM of RNase E and 100 nM of catalytically inactive PNPase in 20 mM tris-HCl pH 7.9, 100 mM NaCl, 10 mM MgCl_2_, 10 μM ZnCl_2_, 0.5% glycerol, 0.01% IGEPAL, and 1 mM DTT for 1 hour at 37°C in a 10 µL reaction. Reactions were stopped by adding 1X of RNA loading dye (ThermoScientific) then heated for 10 minutes at 70°C. Samples were run on a 15% TBE-urea PAGE gel and stained with 40 mL of 1X TBE with 1X Sybr™ Gold (Invitrogen) on a rocker for 10 minutes then imaged with an Azure 600 imager at 302 nm excitation. ImageJ with the FIJI plugin was used to quantify band intensities, which were normalized to a mock reaction with no enzymes.

To test for stimulation of PNPase by RhlE1 (Supplemental Figure 9C), this protocol was modified using a final concentration of 949 nM of the same RNA hairpin (Supplemental Table 3) and 138 nM RhlE1 (N-terminal tags) and 95 nM PNPase in 20 mM tris-HCl pH 6.5, 100 mM NaCl, 5 mM sodium phosphate monobasic, 100 mM MgCl_2_, 0.5% glycerol, 0.01% IGEPAL, 1 mM DTT, and 1 mM ATP in a final volume of 10 µL. Samples were incubated at 37°C for 75 minutes then stopped and imaged as described above.

A 29 nt FAM-labeled RNA derived from the *M. smegmatis atpB* sequence and reported in (54) was purchased from Integrated DNA Technologies and used in cleavage assays with RNase E, RNase J, and PNPase (Figure 3A, Supplemental Figures 3 and 12A, D-E Supplemental Table 3). RNase E and RNase J reaction used the same buffer as described in the first paragraph of this section, and PNPase reactions used the same buffer as described in the second paragraph of this section. 10 µL reactions contained 80 ng of enzymes (final concentrations 101 nM RNase E, 95 nM PNPase, 127 nM RNase J, and 138 nM RhlE1 (N-terminal tags)) and 200 nM RNA and were incubated at 37°C degrees for 1 hour, then stopped by adding 10 µL loading dye (25 mM EDTA pH 8.0 in formamide), heated for 10 minutes at 70°C, run on a 20% urea-PAGE gel, and imaged on an Azure 600 imager using the Cy3 channel (524 nm excitation).

### RNase E Kinetics

The 29 nt FAM-labeled RNA containing exactly one RNase E cleavage site described in the previous section was used for kinetics experiments. For Figure 3, 10 µL reactions contained 10 nM RNase E and 100, 300, 1000, or 2000 nM RNA in buffer composed of 20 mM tris-HCl pH 7.9, 100 mM NaCl, 10 mM MgCl_2_, 10 μM ZnCl_2_, 0.5% glycerol, 0.01% IGEPAL, and 1 mM DTT. Reactions were incubated at 37°C in a thermocycler and stopped at 5-minute intervals by addition of 10 µL loading dye (25 mM EDTA pH 8.0 in formamide) then heated at 70°C for 10 minutes. Samples were run on a 20% urea-PAGE gel then imaged on an Azure 600 imager using the Cy3 channel (524 nm excitation). In FIJI, a box was drawn around each band and an area within the same lane of the gel with no signal. The integrated density was measured and blank subtracted for each lane. A standard curve was run on each gel to calculate the amount of product, and initial rate was calculated by linear fitting of a graph of product versus time with a minimum R^2^ value of 0.8. At least 4 timepoints were used for initial rate calculations, and at least 2 replicates of each initial rate were calculated. Initial rates were plotted versus substrate concentration and were fit using the nonlinear regression K_cat_ model in GraphPad Prism.

Experiments shown in Supplemental Figure 3 were done as above with the following exceptions. The RNase E concentration was 101 nM and the RNA concentrations were 150, 300, 445, 590, 715, 1410, 3020 nM. Reactions were stopped every 30 seconds for 4 minutes, except for reactions with 3020 nM RNA which were stopped every 2 minutes for 16 minutes. Substrate band intensities were quantified using ImageJ with the FIJI plugin and normalized to a mock reaction containing no enzyme run on the same gel. Initial rates were calculated by using the slope of the line of the amount of substrate remaining versus time and were averaged for three replicates. Because the substrate concentrations used were not all substantially higher than the enzyme concentration, we report the results as apparent kinetic parameters.

### Electromobility Shift Assays

HiScribe® T7 High Yield RNA Synthesis Kit (New England Biolabs) kit was used with a 20 µL reaction containing 7.5 µL of NTP + Buffer mix, 1.5 µL of T7 RNA Polymerase, 250 ng of the mCherry DNA template containing a T7 promoter, and 3 µL of fluorescein-12-UTP (Sigma) and incubated overnight at 37°C. 2 µL of DNase I (New England Biolabs) and 30 µL of water were added to the reactions and incubated at 37°C for 15 minutes. RNA Clean and Concentrator-5 (Zymo) was used according to the manufacturer’s instructions with an elution volume of 11 µL.

A final concentration of 50 nM RNA (Figure 3C-D, Supplemental Figure 5, Supplemental Table 3) was mixed with the indicated concentration of catalytically inactive RNase E in 20 mM tris-HCl pH 7.9, 100 mM NaCl, 10 mM MgCl_2_, 10 μM ZnCl_2_, 0.5% glycerol, 0.01% IGEPAL, and 1 mM DTT at 37°C for 15 minutes in a 10 µL reaction. Glycerol was added to a final concentration of 25% for loading the gel. Reactions were run on a 1% agarose gel at 4°C and imaged on an Azure 600 imager Cy3 channel (524 nm excitation). Band intensities were quantified using ImageJ with the FIJI plugin. A box was drawn around the unbound band and an area of the gel with no signal.

The integrated density was measured and blank subtracted for each lane. The unbound fraction was calculated by diving the unbound signal in lane by the 0 nM control. Due to the many multimerization states of RNase E, the fraction bound was calculated performing one minus the unbound fraction, which was then plotted versus the RNase E concentration and fit with a one site - specific binding nonlinear model in GraphPad Prism.

### Coimmunoprecipitations with RNase E

For PNPase and RhlE1 (C-terminal tag), 43 nM catalytically inactive RNase E was mixed with 86 nM catalytically active degradosome enzymes in a final volume of 1 mL with 20 mM tris-HCl pH 7.9, 100 mM NaCl, 5% glycerol, 0.1 mM DTT, 0.01% IGEPAL, 10 mM MgCl_2_, and 10 µM ZnCl_2_ and preincubated with end-over-end rotation at 37°C for 1 hour. For RNase J, the same protocol was used with 21.5 nM catalytically inactive RNase E and 43 nM catalytically active RNase J. 20 µL of anti-FLAG magnetic beads (Sigma) were added and incubated at 37°C for an additional 1 hour. Supernatant was removed and beads were washed 3 times with 150 µL of TBS then eluted with 75 µL of 0.1 M glycine-HCl pH 2.8 and immediately run on SDS-PAGE, loading one tenth the fraction of mock reactions compared to the elutions. Western blots were run as described above, using the same protocol for anti-FLAG antibody and 0.33 µg/mL anti-HA antibody (Biolegend, 901533). ImageJ with the FIJI plugin was used to quantify band intensities.

For coimmunoprecipitations with benzonase (Supplementary Figure 8B), 10 U of benzonase (Millipore) was used in each reaction.

### Coimmunoprecipitation without RNase E

100 ng/µL catalytically active degradosome enzymes (PNPase, RhlE1 (N-terminal tags), or RNase J) in a final volume of 70 µL were combined in 20 mM tris-HCl pH 7.9, 100 mM NaCl, 5% glycerol, 0.1 mM DTT, 0.01% IGEPAL, 10 mM MgCl_2_, and 10 µM ZnCl_2_ and preincubated with end-over-end rotation at 37°C for 1 hour. 12 µL of anti-HA magnetic beads (Sigma) were added and incubated at 37°C for an additional 1 hour. The supernatant was removed and the beads were washed 3 times with TBS then eluted with 30 µL of 0.1 M glycine-HCl pH 2.8 and immediately run on SDS-PAGE, loading an equal fraction of mock reaction and elution. SDS-PAGE gels were stained with Bio-Safe Coomassie G-250 Stain (BioRad) and imaged in an Azure 600 imager. Three-way coimmunopreciptations were done in the same way with 2 µg/µL each protein in a final volume of 70 µL.

### *In vitro* Condensate Formation Microscopy

After size-exclusion chromatography, proteins were dialyzed against buffer containing 20 mM Tris pH 7.4, 150 mM NaCl, 2% glycerol, and 0.1 mM DTT, concentrated to 100 µM, and stored at −80 °C. *E. coli* total RNA was prepared at a stock concentration of 400 ng/µL (in NF H_2_0). Phase separation reactions (10 µL total volume) were assembled by mixing 1 µL protein stock, 1 µL RNA stock, 2 µL of 5X phase separation buffer (90 mM Tris pH 7.4, 500 mM NaCl, 5 mM DTT, 5 mM MgCl₂), and 6 µL nuclease-free water. The final reaction conditions were 10 µM protein, 40ng/uL RNA, 20 mM Tris pH 7.4, 115 mM NaCl, 1 mM DTT, 1 mM MgCl₂, 0.2% glycerol. For reactions containing 10 mM MgCl₂, 50 mM MgCl₂ was used in the 5X buffer. Reactions were incubated at room temperature for 30 min prior to imaging. The entire 10 µL was then spotted on a slide and covered with a coverslip before imaging with a ThermoScientific EVOS microscope with 100X magnification..

The number and diameter of condensates were quantified using the Analyze Particles function in ImageJ. Briefly, three images per protein construct were processed by applying a particle size filter of 0.2 µm to infinity and a circularity range of 0.01 – 1.0. The resulting condensate counts and diameters were then plotted using GraphPad Prism and ordinary one-way ANOVA with Sidak’s multiple comparisons test was used for significance testing. For constructs that did not show detectable condensates, both condensate number and diameter were assigned a value of zero for the purpose of data visualization.

### Phosphate Assay

EnzChek Phosphate Assay Kit (Invitrogen) was scaled down to a final volume of 200 µL in a 96 well plate and read in a Victor Nivo® (Perkin Elmer) multimode plate reader using VICTOR Nivo Control Software® 4.5.0. Reactions containing 50 mM tris-HCl (pH 7.5), 1 mM MgCl_2_, 0.1 mM sodium azide, 200 mM MESG, 0.2 U purine nucleoside phosphorylase, 0.01 µM catalytically active RhlE1 (C-terminal tags), 0.1 µM catalytically inactive PNPase or catalytically inactive RNase E, 1 µM hairpin RNA (Figure 5G, Supplemental Figure 9A, Supplemental Table 3) were preincubated for 10 minutes at room temperature then ATP was added to a final concentration of 2 mM and immediately read on the plate reader every 3 minutes for 1 hour (PNPase) or two hours (RNase E). Absorbance values were normalized to time 0 then converted to phosphate concentration using a standard curve made with KH_2_PO_4_.

### SEC-HPLC

A Nexera Lite Inert High Performance Liquid Chromatography System (Shimadzu) was used with a TOSOH Bioscience G3000SWXL column (7.8 mm diameter, 30 cm length). A program with an isocratic flow of 1 mL/minute with a buffer containing 20 mM Tris-HCl pH 7.9, 100 mM NaCl, 5% glycerol, 0.1 mM DTT, 10 mM MgCl_2_, 10 µM ZnCl_2_, and 0.01% IGEPAL was utilized. Molecular weight size standard (Sigma 69385) was run, and 20 µL of 2.6 µM of each catalytically inactive RNase E variant was injected. UV detection was monitored at 280 nm, and LabSolutions Version 6.117 was used for data collection and analysis.

## RESULTS

### Mycobacterial RNase E has two intrinsically disordered regions

In the organisms where RNase E has been best studied, such as *E. coli* (15,16) and *C. crescentus* (17), this enzyme has an intrinsically disordered region (IDR) that mediates noncatalytic functions including interaction with other proteins and subcellular localization. Mycobacterial RNase E has two IDRs: one on the N-terminus and one on the C-terminus, flanking the catalytic domain (reviewed in (55)). To define the boundaries of the IDRs in *M. smegmatis* RNase E, we used IUPred2 and AlphaFold3 to predict regions of disorder (Figure 1A and B respectively). Multiple sequence alignments in ClustalOmega revealed the expected conservation of this domain architecture in the Actinobacteria, in contrast to the Proteobacteria (Figure 1A and Supplemental Figure 1) (15). The IDRs are quite large, at ∼330 and ∼214 amino acids long in *M. smegmatis* and similar sizes in *M. tuberculosis*. While the IDR sequences are not well conserved across mycobacteria, they consistently include acidic and arginine-rich regions (Supplementary Figure 1). To our knowledge, the Actinobacteria is the only class whose RNase E contains both N- and C-terminal IDRs, so we sought to learn the physiological importance of these regions. To determine the essentiality of the IDRs, we generated an *M. smegmatis* strain expressing full-length RNase E from a plasmid integrated at the L5 phage integration site and deleted *rne* (*MSMEG_4626*) from its native locus. We then exchanged the L5 plasmid for variants encoding truncated RNase E and marked with a different antibiotic resistance gene. All variants had N-terminal FLAG-tags and were expressed from their native promoters with native 5’ UTRs. An N-terminal IDR deletion (retaining AA 331-1037), a C-terminal IDR deletion (retaining AA 2-824), and a N- and C-terminal IDR double deletion (retaining AA 331-824) all supported bacterial survival. In contrast, we were unable to obtain a strain expressing RNase E with a deletion of the first 375 amino acids as its sole copy, suggesting that the essential catalytic domain (7) was compromised (Figure 1C).

**Figure 1.**
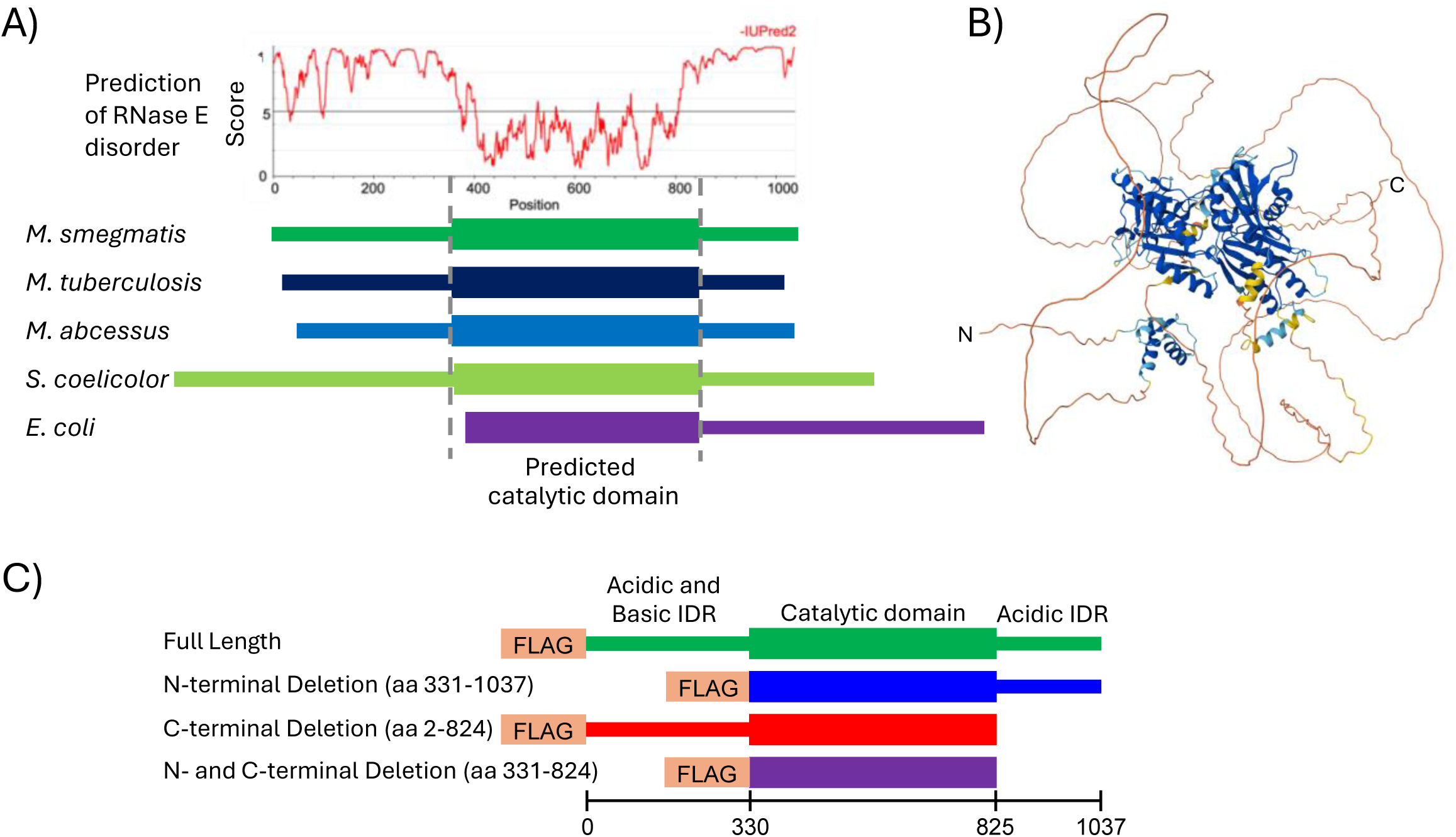
Mycobacterial RNase E has two intrinsically disordered regions. **A)** IUPred2 disorder prediction of *M. smegmatis* RNase E and schematic of multiple sequence alignment (ClustalOmega) showing the location, relative size, and position of RNase E IDRs in the indicated bacteria. **B)** AphaFold3 structure prediction for *M. smegmatis* RNase E. Blue indicates areas of high confidence and red/yellow indicate areas of low confidence. pTM = 0.49. **C)** Schematic of *M. smegmatis* RNase E IDR deletion mutants used in this study.

### The N-terminal IDR of RNase E impacts growth and mRNA degradation rates

To begin to understand the physiological roles of the RNase E IDRs, we investigated the phenotypic consequences of deleting them. First, we performed western blotting to assess the relative amounts of RNase E in the IDR deletion strains and confirmed that each truncated version had abundance similar to, or higher than, the full-length (Figure 2A-B). Therefore, phenotypes we observe with the mutant strains are not due to protein instability upon IDR deletion. We found that deleting the N-terminal IDR resulted in significantly shorter cells compared to strains with full-length RNase E and the C-terminal IDR deletion (Figure 2C). Additionally, the cell size distributions of the N-terminal IDR deletion and N- and C-terminal IDR double deletion strains were broader than those of the strains with full-length or C-terminal deleted versions. This suggests that deleting the N-terminal IDR of RNase E causes a cell growth or division defect. Additionally, the strains with RNase E N-terminal IDR deletions had growth defects observed by optical density (Figure 2D) and colony forming units (CFUs) (Figure 2E).

**Figure 2.**
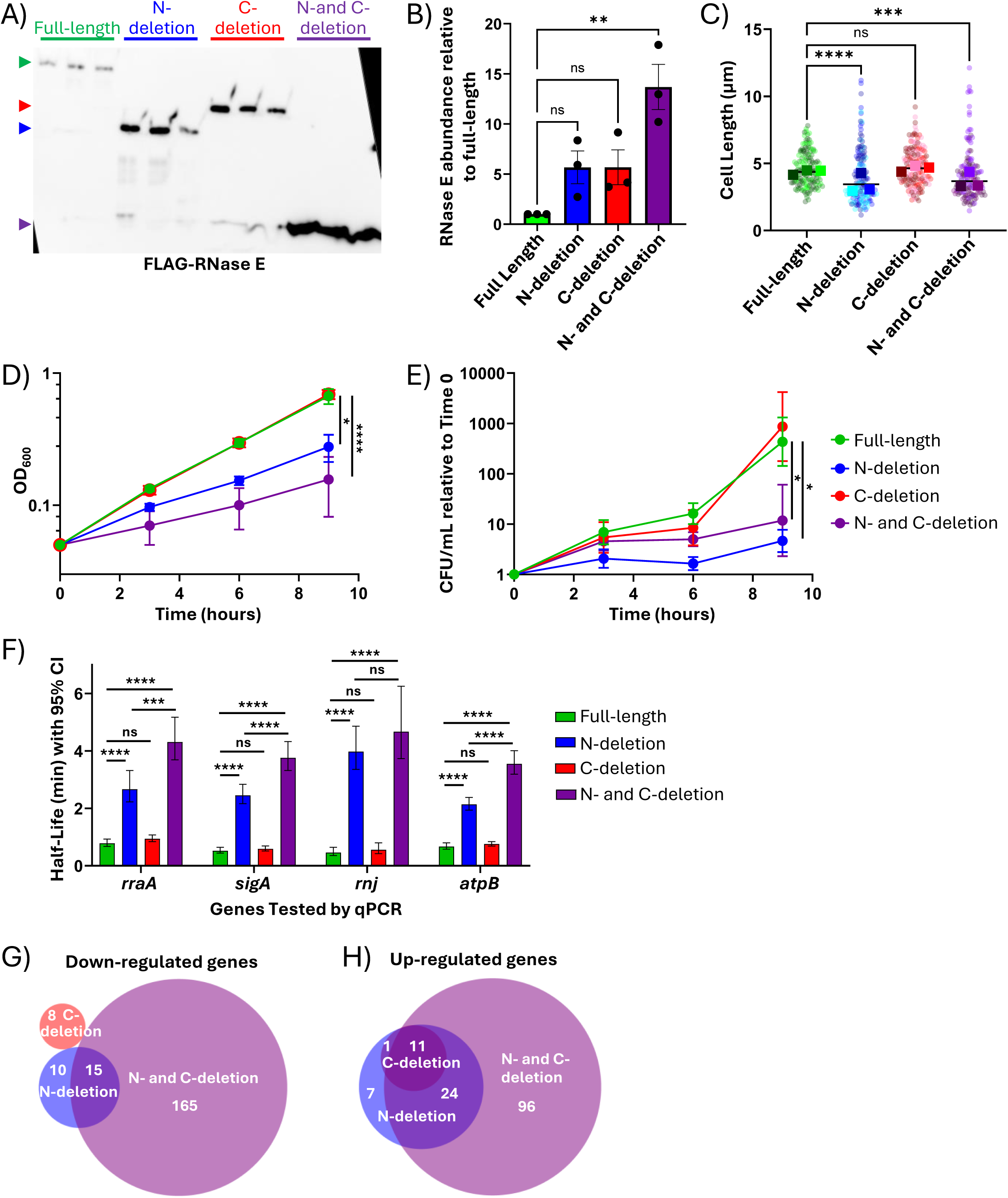
Deletion of the two IDRs of RNase E in *M. smegmatis* has differential impacts on cell size, growth rate, mRNA degradation, and gene expression. All experiments were done with *M. smegmatis* harboring the RNase E variants shown in Figure 1C as sole copies. **A)** Western blot of FLAG-tagged RNase variants in biological triplicate. **B)** The relative abundance of RNase E was calculated from the band intensities in A). The average of three biological replicates was reported and a one-way ANOVA with Dunnett’s multiple comparisons test was used. **C)** Cells were imaged on a spinning disk microscope, and cell length was measured in ImageJ with the FIJI plugin. Circle colors represent biological replicates, and squares represent medians of each biological replicate. Black lines represent the means of the medians. IDR mutants were compared to the strain with full-length RNase E by one-way ANOVA with Kruskal-Wallis multiple comparisons test. **D-E)** Growth was measured by D) optical density and E) colony forming units. Points represent averages of biological triplicate strains. Linear regression was used for significance testing. **F)** RNA half-lives of selected genes were measured by blocking transcription with 150 µg/mL rifampicin and measuring RNA abundance at several timepoints by quantitative PCR. Error bars denote 95% confidence intervals. Half-lives were compared by linear regression. **G-H)** RNAseq was used to compare the steady-state transcriptomes of strains expressing the indicated mutants to that expressing full-length RNase E. Genes were classified as differentially expressed if their log_2_ fold-change was <-1 (downregulated) or >1 (upregulated) in the mutants compared to full-length, with adjusted *p* <0.05. In A-F), ns *p* > 0.05, * *p* < 0.05, * *p* < 0.01, *** *p* ≤ 0.001, **** *p* ≤ 0.0001

To determine if the IDR deletions affected mRNA degradation, we measured the half-lives of the transcripts of four arbitrarily selected genes (*atpB*, *rnj*, *rraA*, and *sigA*). Deleting the N-terminal IDR caused an increase in RNA half-life for all four transcripts, indicating that mRNA was degraded less efficiently. This effect was exacerbated when the C-terminal IDR was additionally deleted (Figure 2F), but deletion of the C-terminal IDR alone did not have a detectable impact. This suggests that the two IDRs have partially redundant functions that are required for normal mRNA degradation rates, while the N-terminal IDR also has a unique function.

To globally assess the impact of the RNase E IDRs on the steady-state transcriptome, we performed RNAseq to identify genes with differentially abundant transcripts. It is important to note that these changes in steady-state transcript abundance likely reflect the combination of two types of effects: (1) differences in degradation rate (direct impacts of perturbed RNase E function) and (2) differences in transcription rate resulting from cellular adaptation to the perturbed RNase E function. Relatively small numbers of genes were differentially expressed in the N-terminal and C-terminal individual IDR deletion strains (68 and 20 respectively), but the double deletion strain showed a large set of differentially expressed genes (311 genes; Figure 2G-H; Supplemental Table 6). This implies that there is functional redundancy between the two IDRs. The proportions of up- and down-regulated genes were similar for all three strains, suggesting that many of the expression changes were due to regulatory responses rather than passive effects of slower mRNA degradation. There were no annotated GO or KEGG pathways enriched in the differentially expressed genes for any of the three strains. However, some pathways were identifiable among differentially expressed genes. Several mycofactocin biosynthesis genes were differentially expressed in the double-IDR deletion strain with similar trends in the N-terminal IDR single deletion strain. This pathway may be involved in redox balance (56). Eleven genes in the ESX-1 type VII secretion system locus were upregulated in the double-IDR deletion strain, of which four were upregulated in the N-terminal IDR single deletion strain. The ESX-1 secretion system is involved in virulence in *M. tuberculosis* and conjugation in *M. smegmatis* (57–59). There were some expression differences in genes associated with DNA replication; *MSMEG_6892*, encoding the replicative DNA helicase, was upregulated in both the N-terminal IDR single deletion strain and the double IDR deletion strain, and *MSMEG_2416*, encoding a septation-associated protein (57,59), was upregulated in the double IDR deletion strain. Additionally, four ribonucleoside reductase genes were upregulated in the double-IDR deletion strain (of which one was also upregulated in the N-terminal IDR single deletion strain) (Supplemental Table 7). The only possible RNase differentially expressed was an annotated endoribonuclease L-psp family protein, which was downregulated in the double-IDR deletion strain. A gene annotated as encoding a ribonuclease inhibitor, *MSMEG_0209*, was also downregulated. The functions of these proteins have not been studied, and the annotations should therefore be treated with caution. None of the four genes shown to have altered degradation kinetics in Figure 2F were differentially expressed, indicating that their slower degradation in the IDR deletion strains was likely compensated by correspondingly slower transcription, as we have previously reported (10).

### The RNase E IDRs contribute to RNA binding but are dispensable for multimerization and catalysis in vitro

Since we uncovered several phenotypes associated with deleting the IDRs of RNase E, we wanted to understand the biochemical mechanism responsible. We recombinantly overexpressed N-terminal 6x-His and FLAG tagged *M. smegmatis* RNase E and the IDR deletion mutants and purified them by immobilized metal affinity followed by size exclusion chromatography. We purified catalytically inactive versions of each RNase E variant (D694R and D738R (10,60) in the full-length protein, and the equivalent residues of each variant) and tested their cleavage activities in parallel to optimize purification procedures to prevent co-purification of *E. coli* RNases. Representative protein preparations are shown in Supplemental Figure 2.

We directly tested the impact of the RNase E IDRs on catalysis by measuring the kinetics of the cleavage of a short 5’ FAM-labeled 29 nt RNA. This substrate was derived from the *atpB* gene and was previously shown to have exactly one RNase E cleavage site (10,54). The kinetic parameters of full-length and double-IDR deleted RNase E were statistically indistinguishable (Figure 3A-B). Preliminary experiments with the single IDR deletions likewise did not suggest differences in kinetic parameters compared to full-length RNase E (Supplemental Figure 3). This indicates that the IDRs do not impact catalysis, which is consistent with the apparent distance of the IDRs from the active site as well as reports that the IDRs of RNase E in other organisms were involved in noncatalytic activities (15–17,40,43).

**Figure 3.**
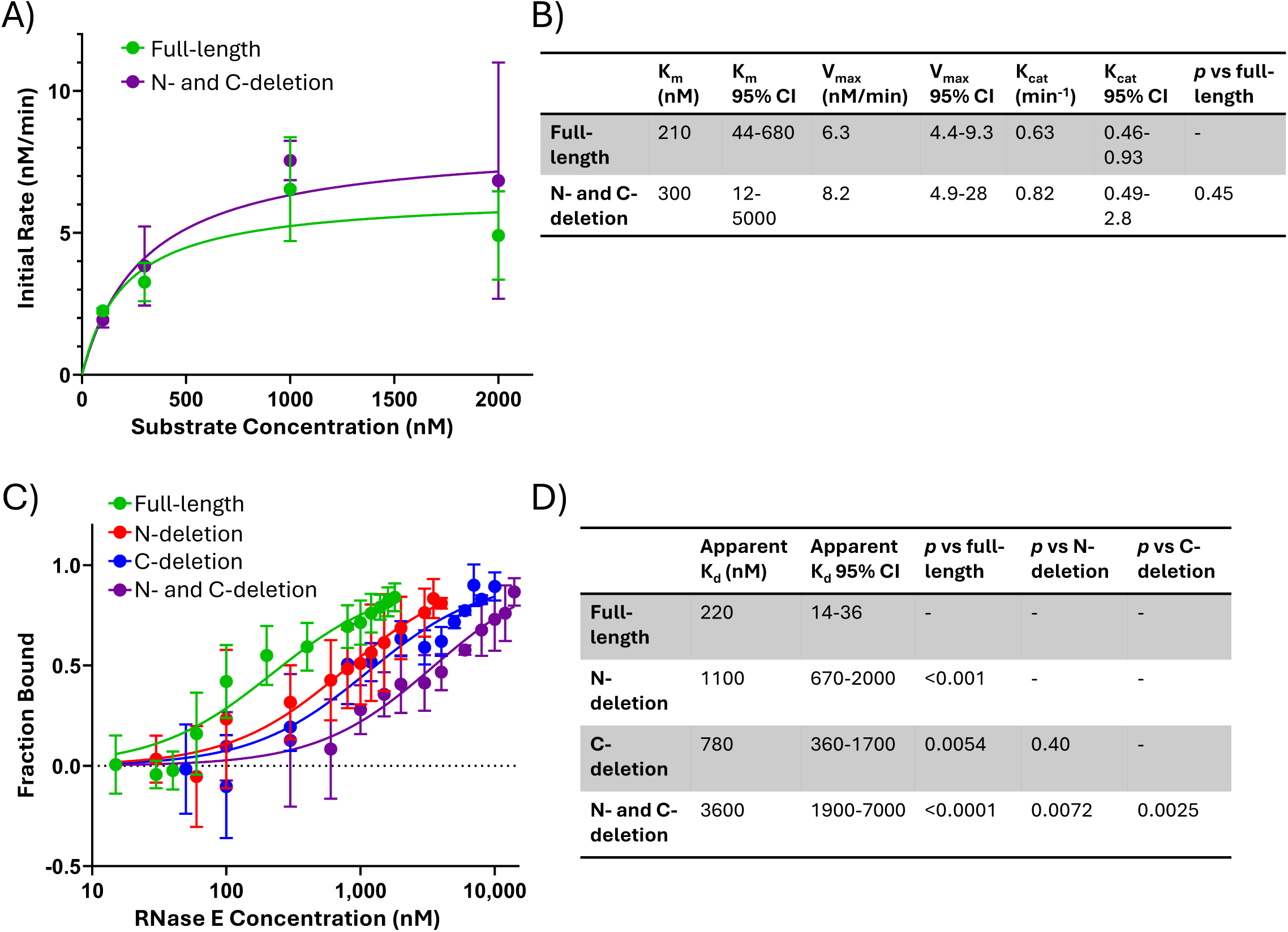
The IDRs of RNase E are dispensable for catalytic activity but have roles in RNA binding. **A)** Kinetic analysis full-length and double-IDR deletion RNase E *in vitro* cleaving a 29 nt FAM-labeled RNA substrate, quantified in **B)**. Initial rates were calculated by simple linear regression of the appearance of product over time, quantified on ImageJ with the FIJI plugin. Points represent the average of two or more replicates. Nonlinear regression with a K_cat_ model was used for determining kinetic constants, and the N- and C-deletion was compared to full-length using the extra sum of squares F-test. **C)** EMSA binding assays with a 721 nt fluorescein-UTP labeled RNA and catalytically inactive versions of the indicated RNase E variants, fit with the one-site specific binding equation quantified in **D)**. The fraction bound was quantified on ImageJ with the FIJI plugin, and each point represents the average of three replicates. Comparisons were made using the extra sum of squares F-test.

In both *E. coli* (12,13) and *M. tuberculosis* (6), RNase E exists as monomers, dimers, and tetramers. To determine if the IDRs impact the multimerization state of *M. smegmatis* RNase E, we performed analytical size-exclusion chromatography. We found monomers, dimers, and tetramers in all IDR deletions, suggesting that the IDRs are not necessary for multimerization (Supplemental Figure 4A-D). Interestingly, the full-length, N-terminal IDR deletion, and double IDR deletion were predominantly in the tetramer state, whereas the C-terminal IDR deletion was primarily in the monomer and dimer states.

Since the IDRs contain arginine-rich regions (Supplemental Figure 1), we hypothesized that they might contribute to RNA binding. To test this, we performed electromobility shift assays (EMSAs) with a 721 nt RNA (Figure 3C-D and Supplemental Figure 5). We reasoned that an RNA of this length would allow for simultaneous binding of multiple sites on RNase E (e.g. the catalytic domain and both IDRs). Due to the three possible RNase E multimerization states as well as the likelihood of multiple RNA binding sites within each protomer, we report apparent K_d_ values that likely reflect the combined affinities of multiple binding sites and modes. Full-length RNase E bound RNA with higher affinity than both the N-terminal and the C-terminal IDR deletions, and the double IDR deletion bound RNA with lower affinity than the single deletions (Figure 3C-D and Supplemental Figure 5). The N-terminal and C-terminal IDR deletions appeared to bind RNA with similar affinities (Figure 3C-D), consistent with each contributing similarly to the overall RNA binding properties of the full-length protein.

### The RNase E IDRs play minor roles in formation of condensate-like bodies

Broadly, bacteria are thought to form biomolecular condensates to quickly bring enzymes and substrates together, allow reactions to happen, then disassemble to allow for rapid responses to environmental changes (reviewed in (18,19)). In *C. crescentus*, the RNase E IDR has an established role in formation of condensates (17,20–22). To determine if *M. smegmatis* RNase E forms condensate-like bodies, we fused RNase E to mCherry to observe its subcellular localization. The RNase E-mCherry fusion was confirmed to be functional because cells were able to grow with this as the only copy of RNase E, an essential protein (7), when the native copy had been deleted. We observed regions of increased RNase E-mCherry fluorescence signal that we will refer to as clusters, and showed that these clusters are dynamic by treating the cells with rifampicin. Clusters were quantified by drawing a line through the long axis of each cell and counting the number of times the fluorescence signal intensity passed above and below the mean as shown in Supplemental Figure 6A. This value was normalized by cell length, as mycobacteria display notable heterogeneity in cell length during log-phase growth. The number of clusters per micron decreased following 30 minutes of rifampicin treatment (Figure 4A-B), consistent with our previous report with an RNase E-mScarlet fusion (61). Since rifampicin blocks transcription, this result suggests that as mRNA levels globally decrease following rifampicin treatment (62), RNase E condensate-like bodies are disassembled. The RNase E clusters did not appear to be associated with the cell membrane, consistent with the lack of a membrane-targeting sequence and with a previous report (63).

**Figure 4.**
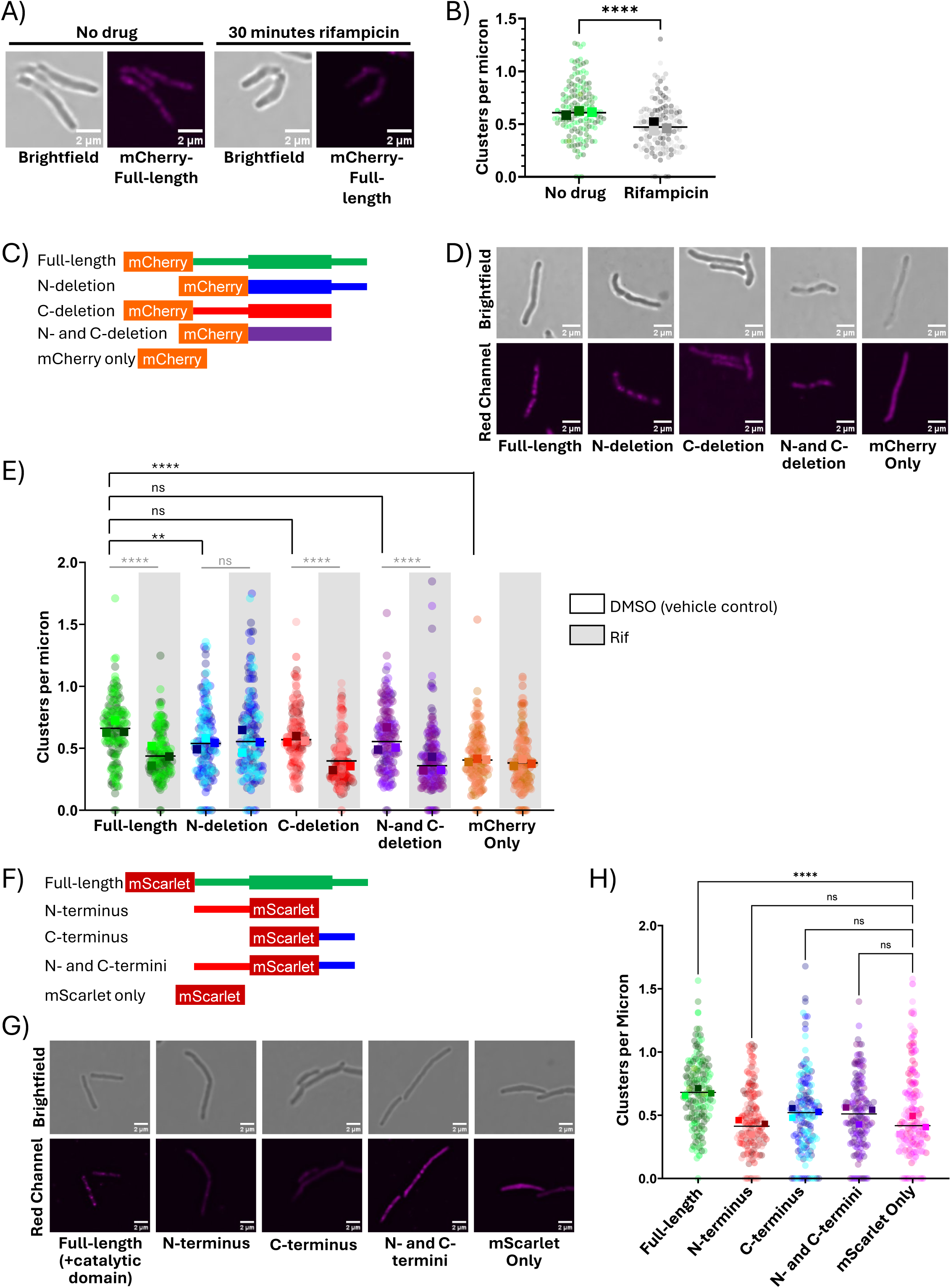
RNase E’s IDRs play minor roles in formation of condensate-like bodies *in vivo*. *M. smegmatis* strains harboring the indicated constructs were imaged by fluorescence microscopy. In A-E, the indicated RNase E variants were present as sole copies. In F-H, strains expressed untagged full-length RNase E from its native locus as well as the indicated constructs. Fluorescence clusters per micron were determined by drawing a line longitudinally through each cell and plotting gray value versus distance in ImageJ with the FIJI plugin. Regions where the signal intensity was greater than the mean were counted as clusters, and the number of clusters was normalized by cell length. **A)** Representative images from fluorescence microscopy of cells expressing mCherry-tagged RNase E with or without 100 µg/mL rifampicin treatment for 30 minutes. **B)** Quantification of A), compared by Mann-Whitney test. **C)** Schematic of mCherry-tagged RNase E constructs for which representative images are shown in **D). E)** Clusters per micron were quantified for cells treated with 100 µg/mL rifampicin for 30 minutes and vehicle controls and compared by ANOVA with Kruskal-Wallis multiple comparisons test. Black lines/asterisks denote comparisons among strains in the untreated control condition, while gray lines/asterisks denote comparisons between conditions. **F)** A schematic of constructs used to assess the ability of RNase E IDRs to induce cluster formation by mScarlet. mScarlet-tagged full-length RNase E was used as a positive control. Representative images are shown in **G)** and quantified in **H)**. For panels B, E, and H, the circle colors represent biological replicates, and the squares represent the median of that biological replicate. Black line represents the mean of the medians. ns *p* > 0.05, * *p* ≤ 0.05, ** *p* ≤ 0.01, *** *p* ≤ 0.001, **** *p* ≤ 0.0001.

To determine if the IDRs of RNase E are required for the formation of condensate-like bodies, RNase E with each domain deletion was fused to mCherry and expressed as a sole copy (Figure 4C-E). We observed a small but significant decrease in the number of fluorescence clusters per micron when the N-terminal IDR was deleted, but these cells still had more clusters than those expressing free mCherry in a wildtype background (Figure 4E). None of the other IDR deletions showed statistically significant changes in condensate-like body formation (Figure 4E). We also treated each of the IDR mutant strains with rifampicin for 30 minutes to test if the condensate-like bodies were dynamic in these strains. Strains harboring all versions of RNase E except the N-terminal IDR deletion had decreased clusters per micron following rifampicin treatment, suggesting that they are dynamic (Figure 4E). Collectively, this suggests that the two IDRs are not strictly required for formation of condensate-like bodies but may differentially influence their nature and number in subtle ways.

To determine if the IDRs alone were sufficient to drive formation of condensate-like bodies, we fused the IDRs to mScarlet individually and in combination without the catalytic domain of RNase E (Figure 4F-H). No differences in clusters per micron were observed between mScarlet alone and the mScarlet-IDR fusions (Figure 4H). Together, these data suggest that while the IDRs of RNase E may contribute to condensate-like body formation in the context of the full-length protein, they are not sufficient to mediate formation of these bodies to the extent seen for the full-length protein.

We also tested the ability of purified RNase E to form condensate-like bodies *in vitro*, finding that full-length, N-terminal IDR deletion, and C-terminal IDR deletion variants of RNase E formed condensate-like bodies visible by phase-contrast microscopy to varying extents. In contrast, the double IDR deletion did not under any conditions (Supplemental Figure 7A-B). Surprisingly, the single IDR deletion variants had a greater propensity for condensate-like body formation than the full-length protein; both the N- and C-terminal deletion variants formed condensate-like bodies with and without 10 mM magnesium and RNA, while the full-length version only formed visible condensate-like bodies in the presence of 10 mM magnesium, a metal bound by RNase E for catalysis. The N-terminal IDR deletion protein formed notably more condensate-like bodies than other variants (Supplemental Figure 7A-B), including the full-length protein, and the C-terminal deletion formed irregularly shaped condensate-like bodies, which may suggest that it reaches the equilibrium state more slowly (64). In contrast to previous reports in *C. crescentus* RNase E (17,20–22), we did not find that RNA stimulated the formation of condensate-like bodies (Supplemental Figure 7A-B) by the full-length protein, but RNA did increase the size of the condensate-like bodies formed by the single IDR deletion mutants (Supplemental Figure 7C). Collectively, these data suggest that the IDRs of mycobacterial RNase E can both stimulate and inhibit formation of condensate-like bodies, depending on local conditions.

### Defining the protein-protein interactions of the *M. smegmatis* degradosome-like network *in vitro*

RNase E binds other mRNA degradation proteins via its IDR in *E. coli* (16,31–35,38) and *C. crescentus* (40). Previous work predicted that PNPase, RNase J, and the helicase RhlE may interact with RNase E to form the RNA degradosome in mycobacteria, based on RNase E pull-downs from cell lysates followed by mass spectrometry (11). However, those experiments could not distinguish direct from indirect or RNA-mediated interactions, and did not assess the roles of RNase E’s IDRs. We therefore performed coimmunoprecipitations with purified proteins to probe the direct protein-protein interactions of the degradosome-like network. RNase E was pulled down with its FLAG tag, and the protein partner was western blotted for its hemagglutinin (HA) tag. RNase E was also western blotted for FLAG in each experiment as a control. Western blotting was necessary to obtain clear data in these experiments because RNase E was somewhat unstable during the immunoprecipitation process, resulting in the presence of degradation products of similar sizes as the other protein partners (Supplemental Figure 8A). *M. smegmatis* has two homologs of RhlE (RhlE1 and RhlE2) but *M. tuberculosis* only has one. We chose to test RhlE1 as it has greater sequence similarity to *M. tuberculosis* RhlE.

We identified direct interactions between RNase E and PNPase (Figure 5A-B), between RNase E and RNase J (Figure 5C-D), and between RNase E and RhlE1 (Figure 5E-F). Deletion of the C-terminal IDR of RNase E greatly reduced its ability to pull down PNPase, while deletion of the N-terminal IDR had no detectable impact (Figure 5A-B). Surprisingly, RNase E with a double IDR deletion appeared to pull down more PNPase than that with the C-terminal IDR deletion alone, suggesting that both the catalytic domain and C-terminal IDR of RNase E contribute to PNPase binding. To confirm the interaction was not mediated by RNA potentially co-purified with the proteins, we treated samples with a nuclease during the coimmunoprecipitation and observed the same result (Supplemental Figure 8B).

**Figure 5.**
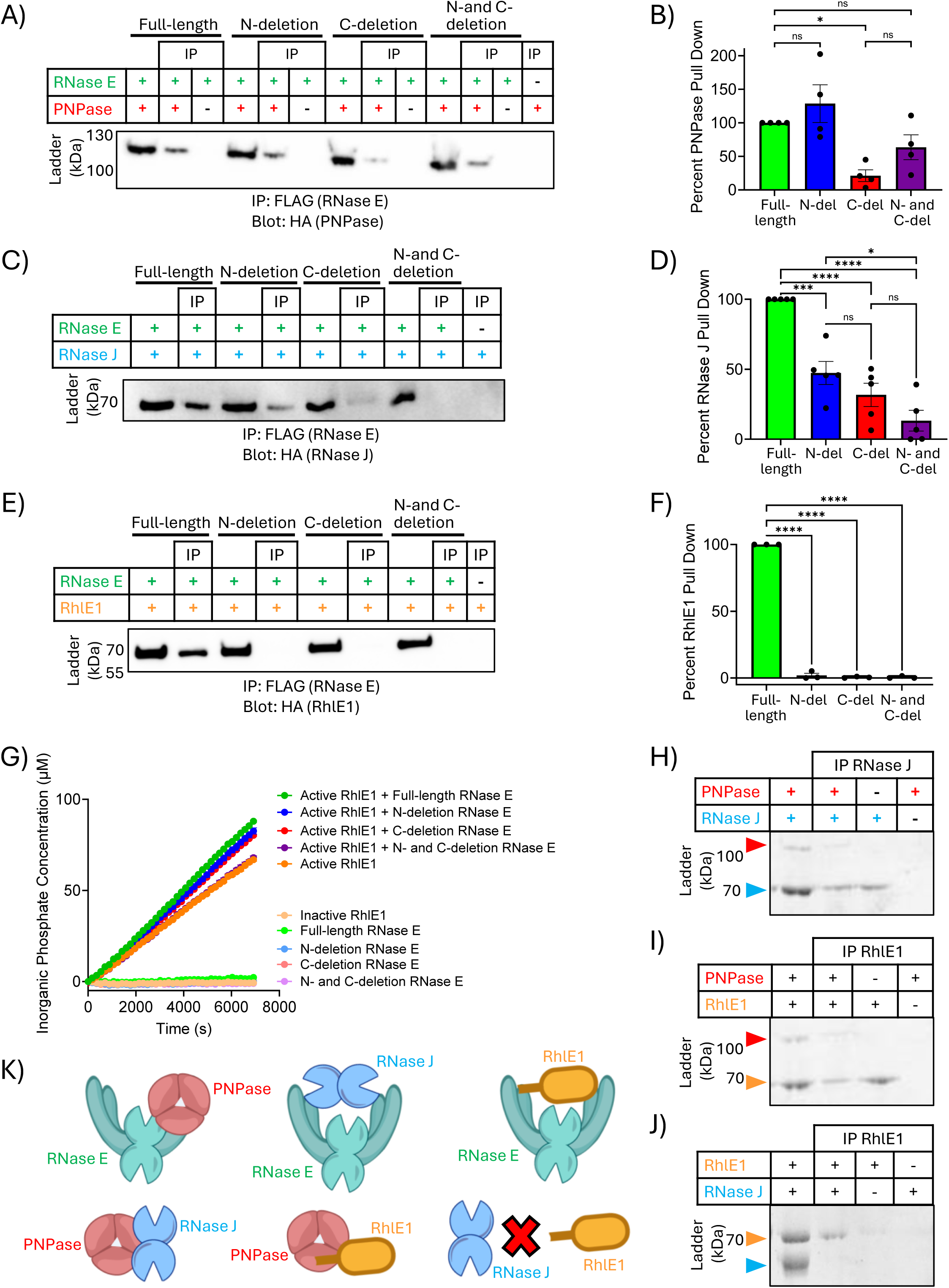
Defining the protein-protein interactions of the degradosome-like network *in vitro*. Representative images of pull downs of FLAG-tagged RNase E and coimmunoprecipitation of HA-tagged **A)** PNPase, **C)** RNase J, and **E)** RhlE1, detected by western blot against HA. In all cases “IP” indicates immunoprecipitation, and samples incubated in buffer without beads were run in parallel for comparison. Pull downs were quantified in **B)**, **D)**, and **F)**, respectively, using ImageJ with the FIJI plugin. Pull down by IDR deletions were expressed as percentages of the full-length pull down. Dots represent replicates, and bars represent averages. A one-way ANOVA was performed with Sidak’s multiple comparison test. **G)** Phosphate production assay of RhlE1 in combination with different versions of catalytically inactive RNase E. Points represent the average of three replicates and simple linear regression was performed. Error bars represent the standard error of the mean for each time point. Representative images of Coomassie-stained SDS-PAGE gels of HA coimmunoprecipitations of **H)** HA-RNase J and untagged PNPase, I) HA-RhlE1and untagged PNPase, and J) HA-tagged RhlE1 and untagged RNase J. **K)** Summary of the verified 2-way protein-protein interactions. ns p > 0.05, * p ≤ 0.05, ** p ≤ 0.01, *** p ≤ 0.001, **** p ≤ 0.0001.

The interaction between RNase J and RNase E was partially reduced for the single IDR deletions and greatly reduced for the double IDR deletion (Figure 5C-D), suggesting that both IDRs participate in the interaction in a roughly additive fashion. In contrast, both IDRs appeared to be required for the interaction between RhlE and RNase E, as RhlE was nearly undetectable in pulldowns when either or both IDRs were deleted (Figure 5E-F).

In *E. coli*, interaction with RNase E causes the RNA helicase RhlB to unwind RNA at a faster rate (16,28,29). To assess if RNase E impacts the activity of RhlE1 in mycobacteria, we utilized a phosphate release assay with a structured hairpin RNA. RhlE1 is an ATP-dependent RNA helicase that produces inorganic phosphate when it unwinds RNA, and we used a catalytically inactive mutant (K55A mutation (65)) as a negative control. With the addition of full-length catalytically inactive RNase E, we observed an increase in the rate of ATP hydrolysis by RhlE1 (Figure 5G) analogous to what was reported for *E. coli*. The N-terminal and C-terminal RNase E IDR single deletions also stimulated RhlE1 but not to the same extent as full-length RNase E, and the double IDR deletion did not stimulate RhlE (Figure 5G). Collectively, these data suggest that both IDRs participate in the interaction between RNase E and RhlE1, and this interaction stimulates RhlE1 activity. Our inability to detect binding between RNase E single-IDR deletions and RhlE1 by immunoprecipitation suggests that the interactions between each IDR and RhlE1 may be transient or weak.

A direct interaction between *M. tuberculosis* PNPase and RNase J was previously reported (11). We wanted to confirm that this interaction also occurred in *M. smegmatis* and to probe additional possible interactions among PNPase, RNase J, and RhlE1. We performed reciprocal coimmunoprecipitations, pulling down HA-tagged versions of each protein and assessing if the other untagged protein coimmunoprecipitated by SDS-PAGE and Coomassie staining. We identified direct interactions between PNPase and RNase J (Figure 5H and Supplemental Figure 8C) as well as between PNPase and RhlE1 (Figure 5I and Supplemental Figure 8D). In both cases, these interactions were observed when either partner protein was pulled down. However, we did not detect an interaction between RNase J and RhlE1 when either of the two proteins was pulled down (Figure 5J and Supplemental Figure 8E).

We propose a model in which these pairs of interactions could occur individually, as we have demonstrated by coimmunoprecipitation *in vitro* (Figure 5K). It is possible that multiple interactions could occur simultaneously as was shown in *E*. *coli* (31,35), but we have no direct evidence for that idea in mycobacteria. To probe multiple simultaneous interactions, we performed a three-component coimmunoprecipitation experiment pulling down HA-tagged RhlE1 in the presence of both PNPase (its direct interaction partner) and RNase J (which could only be pulled down via its interaction with PNPase). While RhlE1 pulled down PNPase as expected, RNase J was not pulled down, indicating that a stable ternary complex was not formed (Supplemental Figure 8F). This suggests that either the interactions of PNPase with RhlE1 and RNase J are mutually exclusive, or the simultaneous interactions are too weak to be identified by pulldown.

Having found that RNase E stimulates RhlE1, we used the same phosphate production assay to test if catalytically inactive PNPase (R431A mutation (8)) enhances RhlE1’s ATPase activity, but we observed no changes (Supplemental Figure 9A). We then used RNA cleavage assays to assess the cleavage activities of protein combinations using the same structured hairpin RNA. We observed no differences in the cleavage activity of RNase E in the presence of catalytically inactive PNPase (Supplemental Figure 9B) or the cleavage activity of PNPase in the presence of RhlE1 (Supplemental Figure 9C).

### Localization of degradosome-like network proteins in vivo

To determine if the degradosome-like network protein-protein interactions identified *in vitro* are reflected in the subcellular localization patterns of these proteins *in vivo*, we performed microscopy and assessed the localization of mCherry-tagged RNase E and partner proteins tagged with green fluorescent protein (GFP). PNPase was tagged with eGFP, and the functionality of the fusion was confirmed using a growth curve with PNPase knockdown using the CRISPRi system (66). PNPase is essential in mycobacteria and causes a growth defect when knocked down, but complementation with a CRISPRi-resistant PNPase-eGFP fusion restored growth to wildtype levels (Supplemental Figure 10A). RNase J was tagged with Dendra2, and its functionality was tested by a rifampicin killing curve because rifampicin causes an increase in killing in an RNase J knockout in *M. smegmatis* (which is notably different from *M. tuberculosis* where knockout of RNase J increases rifampicin tolerance (24)). Complementation of an RNase J deletion strain with the RNase J-Dendra2 fusion restored killing back to wildtype levels (Supplemental Figure 10B). We are not aware of phenotypes for RhlE1 deletion strains in *M. smegmatis*, so we were unable to test its functionality *in vivo*. RhlE1 has a C-terminal IDR, and we fused Dendra2 to the C-terminus of the protein to minimize potential impact to the catalytic domain. In these strains, the sole copy of RNase E was fused to mCherry (with the IDR mutants as in Figure 4C-E), and they were merodiploid for the other degradosome proteins that were fused to GFP.

To quantify the degree to which the proteins were localized in the same space in the cell, the fluorescence intensity in the green channel (PNPase, RNase J, or RhlE1) was plotted versus the signal in the red channel (RNase E), and the Pearson’s correlation coefficient was reported (see examples in Supplemental Figure 11A-E). All three partner proteins (PNPase, RNase J, and RhlE1) had localization patterns that were significantly correlated with that of full-length RNase E (Figure 6). Similar to the *in vitro* immunoprecipitation results, PNPase showed reduced localization with RNase E *in vivo* when the C-terminal IDR of RNase E was deleted (Figure 6A-B), suggesting that its localization in cells is closely tied to its interaction with the C-terminal IDR of RNase E. Unexpectedly, deletion of the N-terminal IDR of RNase E led to a small but statistically significant increase in localization with PNPase, which could potentially be related to the apparently lower dynamicity of the condensate-like bodies formed by this RNase E mutant (Figure 4F). This idea is also consistent with the apparently increased propensity of the N-terminal IDR mutant to form condensate-like bodies *in vitro* (Supplementary Figure 7).

**Figure 6.**
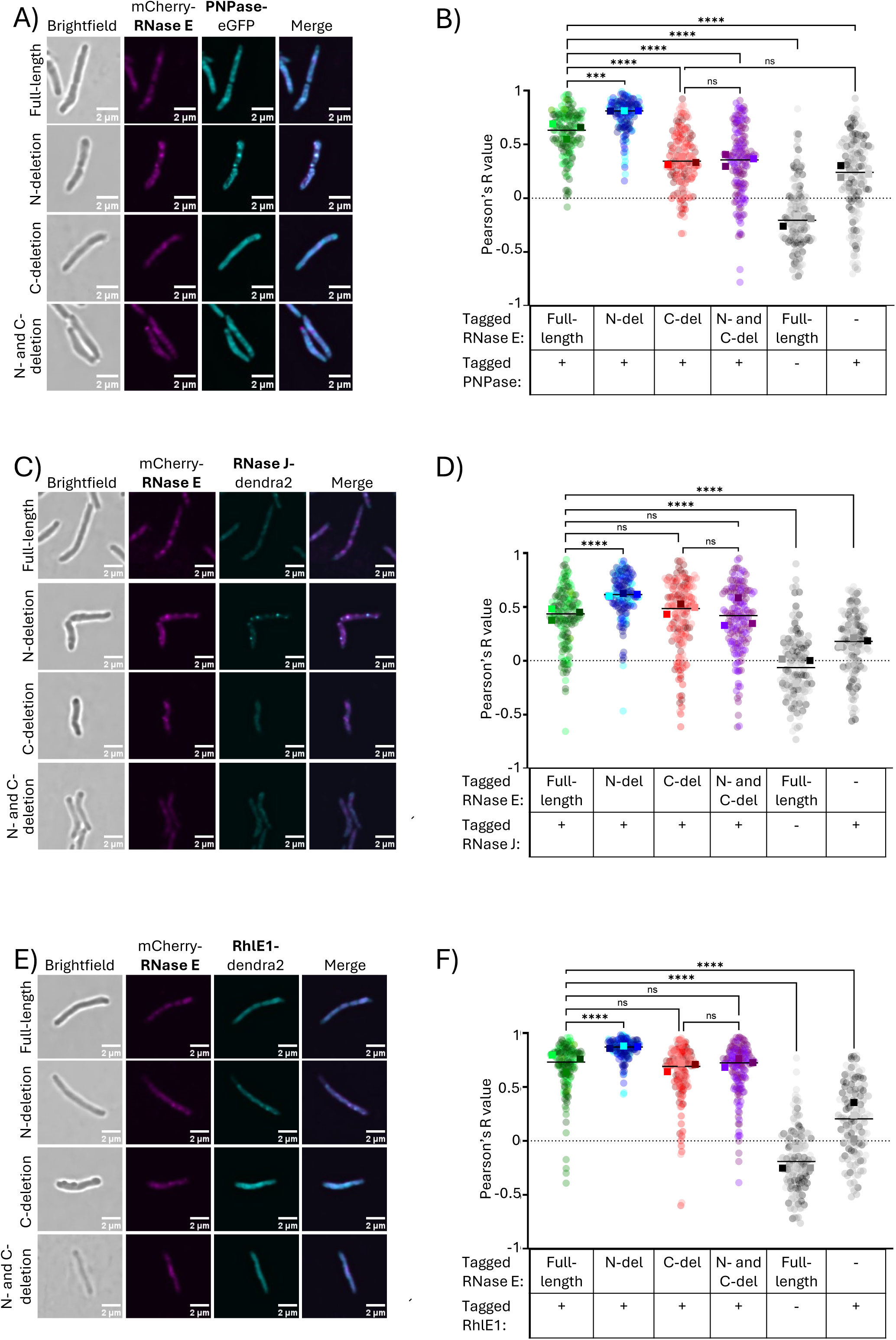
Localization of degradosome-like network proteins *in vivo.* Representative cells from spinning disk microscopy of RNase E-mCherry with IDR deletions and **A)** PNPase-eGFP, **C)** RNase J-dendra2, and **E)** RhlE1-dendra2. Quantification of localization in **B)**, **D)**, and **F)**, respectively, was done by drawing a line through a cell in ImageJ with the FIJI plugin, plotting green versus red channels, and computing the Pearson’s R value. Values at +1 indicate complete correlation, 0 indicates no correlation, and −1 indicates anticorrelation. Circle colors represent biological replicates, and squares represent the median of that biological replicate. Black line represents the mean of the medians. A one-way ANOVA with the Kruskal-Wallis multiple comparisons test was performed. ns p > 0.05, **** p ≤ 0.001 **** p ≤ 0.0001.

RNase J also showed a modest increase in localization with RNase E upon deleting the N-terminal IDR of RNase E, but it localized with the C-terminal IDR deletion and double IDR deletion of RNase E similarly to full-length RNase E (Figure 6C-D). This suggests that the interaction observed *in vitro* between RNase J and the C-terminal IDR of RNase E is not a major contributor to the localization of RNase J in cells.

The impact of the RNase E IDRs on subcellular localization patterns of RhlE1 was very similar to that of RNase J. RhlE1 had a modest increase in localization with RNase E when the N-terminal IDR of RNase E was deleted, but the C-terminal IDR deletion and double IDR deletion of RNase E correlated with RhlE1 to the same extent as full-length RNase E (Figure 6E-F), suggesting that as for RNase J, other factors dictate its localization within cells. RhlE1 itself has an IDR on its C-terminus, so it is possible that this region plays a dominant role in its localization in cells as condensate formation was shown to be conserved among RNA helicases in bacteria (21).

### *In vivo* localization of degradosome-like network proteins changes during carbon starvation stress

Inside the human host, *M. tuberculosis* faces many stressors including hypoxia (67,68). Hypoxia has been shown to dramatically reduce rates of mRNA degradation in mycobacterial cells (3,4) which has been explained as an energy conservation mechanism during periods of stress for the bacteria. Carbon starvation also causes stabilization of mRNA (3), and like hypoxia is a type of energy stress. We hypothesized that weak, transient interactions and localization patterns within the cell were primarily responsible for efficient cleavage rates during log phase growth, rather than fixed interactions or post-translational modifications such as phosphorylation. To investigate this, we first confirmed that the degradosome enzymes RNase E, PNPase, and RNase J cleaved the 29 nt fluorescently labeled oligo from Figure 3A-B (Supplemental Figure 12A) (54). To identify which enzyme was primarily responsible for cleavage of this short RNA in a cell lysate, we utilized the CRISPRi system to knock down PNPase in wildtype and RNase J knockout backgrounds (Supplemental Figure 12B-C). An RNase E knockdown strain was made by replacing the native promoter with an inducible one in the chromosome (10). We then lysed these strains, expecting that lysis would disrupt weak interactions and subcellular localization patterns but not strong interactions or post-translational modifications. We assayed cleavage of the fluorescently labeled RNA by lysates. We found that PNPase was primarily responsible for the cleavage of this short oligo in a lysate (Supplemental Figure 12D and F) and that RNase E knockdown had no impact (Supplemental Figure 12E-F). This suggests that RNase E may have only a minor role in cleavage of short RNAs in cells, consistent with its suggested role in initiation of mRNA degradation. To assess the impact of stress on the ability of PNPase to cleave short RNAs in lysates, we used a 22 nt fluorescently labeled RNA and performed cleavage assays using lysates from cells that had been grown in hypoxia or carbon starvation stress since these conditions are known to stabilize mRNA globally. We found that there were no differences in cleavage rates in lysates from either stress condition compared to log phase (Supplemental Figures 12G-J), consistent with the idea that transient interactions and localization patterns control mRNA cleavage rates in mycobacteria.

Furthermore, we looked at the impact of stress on RNA degradation protein localization in living *M. smegmatis* cells by assessing protein localization with live-cell microscopy using the previously generated fluorescently labeled protein strains from Figure 6. Imaging of hypoxic cells was not technically feasible with our imaging system. Carbon starvation was therefore utilized to assess *in vivo* localization of degradosome proteins in *M. smegmatis* in a condition causing stress-induced mRNA stabilization (3). First, the localization patterns of each protein were individually assessed by counting clusters per micron during log phase growth (carbon rich) and following 24 hours of carbon starvation (Figure 7A, and Supplemental Figure 6A). In all cases, protein localization was impacted by carbon starvation. RNase E, PNPase, and RNase J showed a decrease in the number of clusters per micron. RhlE1 had the opposite trend, showing less clusters than the other three proteins during log phase growth but increased cluster formation during carbon starvation.

**Figure 7.**
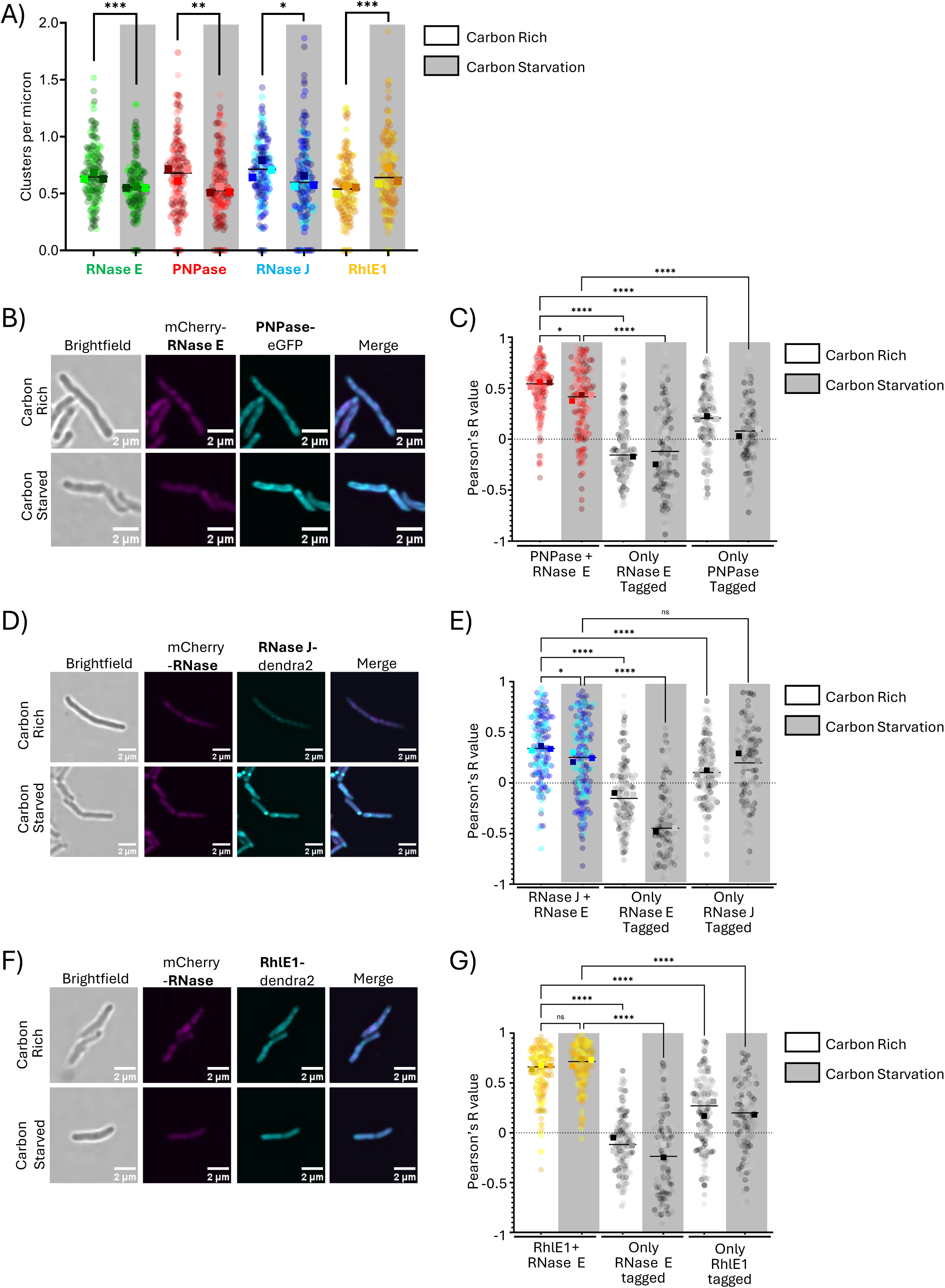
Localization of degradosome-like network proteins *in vivo* during carbon starvation stress. **A)** Single-tagged strains of mCherry-RNase E, PNPase-eGFP, RNase J-dendra2, and RhlE1-dendra2 were imaged by spinning disk microscopy during both carbon rich and carbon starvation conditions. Foci per micron was calculated as in Figure 4B. Representative cells from spinning disk microscopy of RNase E-mCherry with IDR mutants and **B)** PNPase-eGFP, **D)** RNase J-dendra2, and **F)** RhlE1-dendra2 are shown during carbon rich and carbon starvation conditions. Quantification of localization in **C)**, **E)**, and **G)**, respectively, was done as in Figure 6 B, D, and F. The circle colors represent biological replicates, and the squares represent the median of that biological replicate. Black line represents the mean of the medians. A one-way ANOVA with the Kruskal-Wallis multiple comparisons test was performed. ns p > 0.05, **** p ≤ 0.0001.

The correlations of PNPase, RNase J, and RhlE1 localization with RNase E were also assessed in carbon starvation in a similar way as in Figure 6. PNPase (Figure 7B-C) and RNase J (Figure 7D-E) showed small but statistically significant decreases in Pearson’s R values during carbon starvation stress. The changes were likely small due to the proteins spreading out to similar extents leading to more uniform signals across both channels in the cell during carbon starvation. RhlE1 did not show a statistically significant change in its correlation with RNase E (Figure 7F-G). The verified protein-protein interactions among RNA degradation proteins and the changes in protein localization during carbon starvation are summarized in Figure 8.

**Figure 8.**
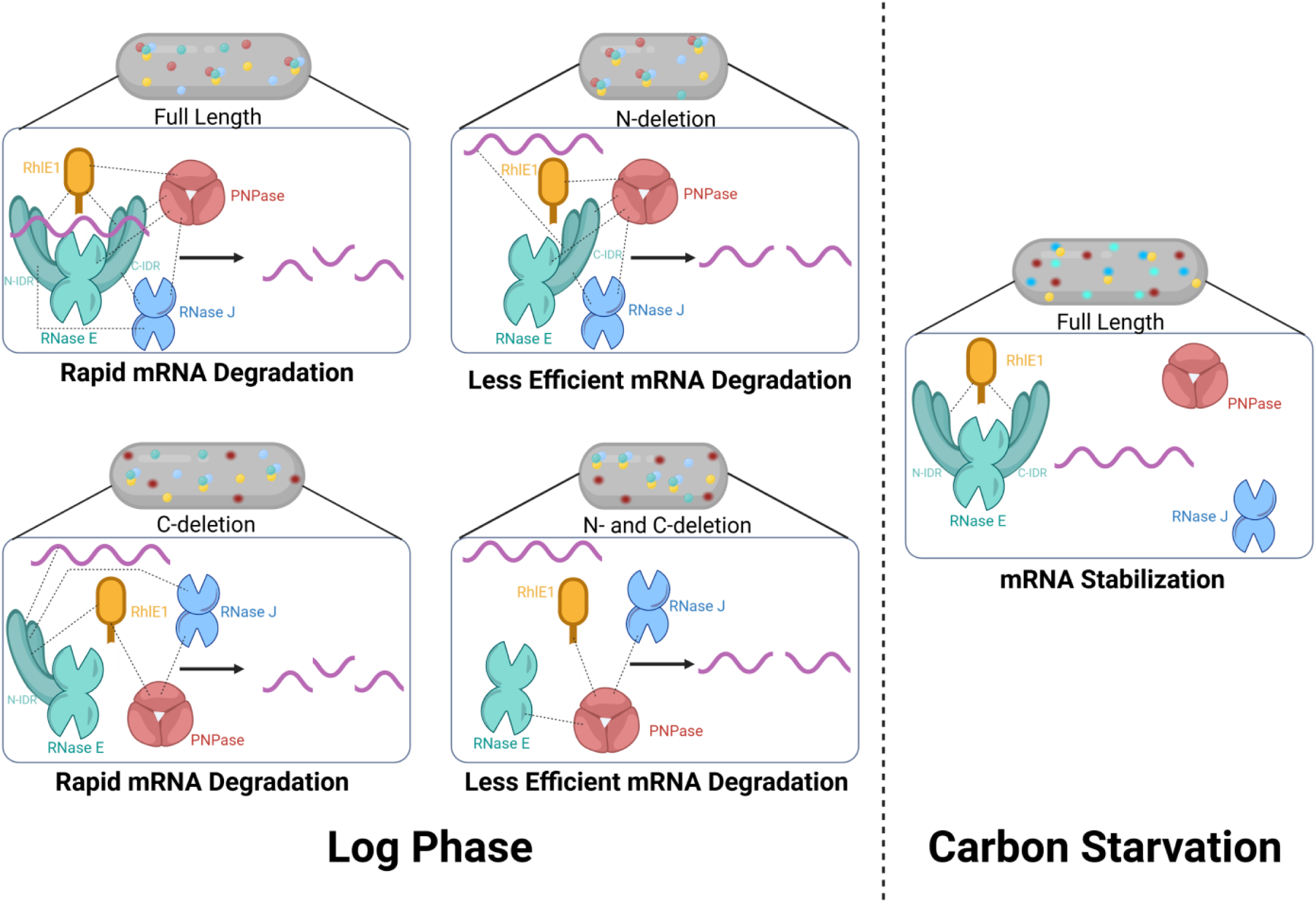
Model of the degradosome-like network in *M. smegmatis* during log phase with RNase E IDR deletion mutants and in carbon starvation. The circles in the cells show the foci identified *in vivo* with teal representing RNase E, blue representing RNase J, red representing PNPase, and yellow representing RhlE1. Fuzzy circles represent diffuse foci. Black dashed lines represent transient interactions with RNase E identified *in vitro*. Created in BioRender. Rapiejko, A. (2026) https://BioRender.com/xvtk9k8.

## DISCUSSION

In *E. coli*, RNase E controls global mRNA decay rates (69–71), and this activity is influenced by the degradosome that is scaffolded by its one IDR (15,16,38). While mycobacterial RNase E was known to have a similarly major role in RNA degradation (10), its interactions with other putative degradosome components were previously unknown. Here, we demonstrate that the two IDRs of mycobacterial RNase E are nonessential in *M. smegmatis* and do not affect catalytic activity on a short oligo substrate. Deleting either IDR resulted in a relatively small number of differentially expressed genes, but the double deletion showed many upregulated and downregulated genes indicating substantial functionally redundancy. Consistent with this, the IDRs both contribute to RNA binding and facilitating interactions with RNase J and the helicase RhlE1 in partially redundant ways. The IDRs also contribute to formation of condensate-like bodies and interaction with PNPase. Furthermore, we have defined many of the direct protein-protein interactions that mediate formation of degradosomes or, more likely, degradosome-like networks in mycobacterial cells. In *E. coli*, the degradosome was defined using techniques including copurification, size exclusion chromatography, yeast-two hybrid assays, and coimmunoprecipitation from lysates without crosslinking (15,16,31,33,34) leading to the conclusion that a stable, stoichiometrically defined degradosome exists in cells. We had to employ more sensitive techniques to probe the degradosome-like network in *M. smegmatis,* suggesting that a less rigid system exists. In our hands, the interactions between degradosome components were too weak to analyze by size-exclusion chromatography. We were also unable to immunoprecipitate degradosomes from *M. smegmatis* or *M. tuberculosis* cell lysates; in the absence of crosslinking, RNase E did not pull down any other proteins in appreciable quantities, and the addition of crosslinking agents caused RNase E to partition into very large, insoluble complexes. Collectively with our *in vivo* localization data, these observations and our *in vitro* data are consistent with the idea of a degradome-like network as has been previously described in *B. subtilis* (reviewed in (45)), with many relatively weak protein-protein and protein-RNA interactions producing a dynamic RNA degradation network.

RNA degradosomes that include both RNase E and RNase J have not been previously described in other bacteria. For the first time, we have identified a direct protein-protein interaction between these two proteins, as well as interactions between each protein and other degradation proteins. RNase E directly interacts with the other three RNA processing and degradation enzymes that we tested, consistent with expectations for an RNase E-scaffolded degradosome similar to those of the Proteobacteria. Previous work in *M. tuberculosis* suggested, but did not directly test, the presence of PNPase-scaffolded degradosomes as well (11), which has been defined in eukaryotic mitochondria by association with the RNA helicase Suv3 (reviewed in (72)). Our *in vitro* results were also consistent with this idea; PNPase directly binds each of the three other enzymes that we tested and may have a hub-like role. RNase J and RhlE1 each bind to two other enzymes, implying a great number of multi-protein complex configurations are therefore possible. However, a complex of more than two different degradosome components in mycobacteria remains hypothetical. We did not detect the only possible three-way interaction that we were able to assay by coimmunoprecipitation in this study (Supplemental Figure 8F).

Cytoplasmic organization of RNA processing and degradation by biomolecular condensates has been found in both prokaryotes and eukaryotes. In *C. crescentus,* RNase E condensates have been shown to stimulate mRNA degradation with the decay process being initiated by an RNase E cleavage event (22). The RNase E IDR was necessary for condensate formation in *C. crescentus* (17), where RNase E can recruit PNPase to condensates to stimulate PNPase activity on cleaved RNAs (20). In mycobacteria, we observed that RNase E’s IDRs were partially redundant and necessary for condensate-like body formation *in vitro* but not *in vivo*. One possible explanation for this discrepancy is that the double IDR deletion formed bodies that were not large enough to be visualized by diffraction-limited microscopy. Another possibility is that the *in vitro* condensate-like bodies are fundamentally different than the *in vivo* ones, possibly because they lack additional components found in cells such as other proteins. *In vivo*, these condensate-like bodies appear to be more like P-bodies in yeast, where disrupting one component does not disrupt the entire condensate (73), and multiple proteins from the body can drive phase-separation (74), furthering the idea of a degradosome-like network held together by a combination of RNA binding, condensate-like bodies, and transient protein-protein interactions. The *in vitro* assay is likely an oversimplification of what is happening in cells, given that we assayed RNase E in the absence of other proteins.

We found that the RNase E’s IDRs contribute to protein-protein interactions, but the importance of these roles in cells differed from what our *in vitro* data suggested. Our *in vitro* data predicted that the C-terminal IDR of RNase E was important for all degradosome interactions. However, the impact of this IDR on protein colocalization *in vivo* was only evident for PNPase. For RNase J and RhlE1, loss of direct protein-protein interactions with RNase E *in vivo* may be masked by other factors that draw the proteins together in condensate-like bodies, such as interactions with RNA. Furthermore, the C-terminal IDR deletion did not affect degradation rates of the transcripts we tested, suggesting that interactions of RNase E with PNPase, RNase J, and RhlE1 are not required for normal mRNA degradation rates. These data contrast with findings that deletion of the C-terminal RNase E IDR reduces degradation efficiency of mRNA in *E. coli* (38) and 5’ UTR cleavage in *C. crescentus* (17).

Surprisingly, we observed increased colocalization of RNase E with the three other degradosome proteins when the N-terminal IDR of RNase E was deleted, despite this mutant having decreased interaction with RNase J and RhlE1 *in vitro*. We also found that the fluorescence clusters of the N-terminal IDR deletion in cells were not disrupted by addition of rifampicin, suggesting that these RNase E clusters are qualitatively different from those of the other variants. Finally, deleting the N-terminal IDR caused less efficient degradation of the mRNAs that we tested, despite causing increased colocalization with other degradosome proteins. Collectively, our data highlight the complexities of the factors that influence mRNA degradation rates in cells; a C-terminal IDR deletion that disrupts protein-protein interactions does not impact mRNA degradation rates, while an N-terminal IDR deletion that appears to increase physical proximity of proteins in cells causes mRNA degradation rates to slow.

Finally, we demonstrated that the subcellular localization patterns of RNA degradation proteins in *M. smegmatis* change in response to carbon starvation, a condition that slows mRNA degradation. This represents a possible mechanism by which mycobacteria may regulate mRNA decay in response to stress. RNA stabilization is a clinically relevant stress response in *M. tuberculosis*, as it occurs during hypoxia (4,65) which occurs inside granulomas (67,68). However, testing the idea that RNase localization controls mRNA degradation is complicated by our finding that these localization patterns are largely robust to mutations that disrupt specific protein-protein interactions, as discussed above.

The results presented here support the IDRs of RNase E playing multiple noncatalytic roles in the cell including RNA binding, condensate-like body formation, and protein-protein interactions. We propose a model in which RNase E forms a degradosome-like network composed of dynamic, transient interactions with other proteins in cells that can be disrupted upon RNA stabilization with a physiologically relevant stressor.

## ACKNOWLEDGEMENTS

We thank members of the Shell lab for helpful discussions, and the UMass Medical School Deep Sequencing core for RNAseq. Graphical abstract created in BioRender. Rapiejko, A. (2026) https://BioRender.com/q1afizi.

## AUTHOR CONTRIBUTIONS

**Abigail R. Rapiejko:** conceptualization, methodology, formal analysis, investigation, writing – original draft, data visualization. **Ying Zhou:** conceptualization, methodology, investigation, data visualization. **Vidhyadhar Nandana:** methodology, investigation, formal analysis. **Junpei Xiao:** formal analysis. **Jared M. Schrader:** methodology, supervision. **Scarlet S. Shell:** conceptualization, writing – review & editing, supervision, project administration, funding acquisition.

## SUPPLEMENTARY DATA

**Supplemental Figures 1-12**

**Supplemental Table 1. *M. smegmatis* strains**

**Supplemental Table 2. Plasmids integrated into *M. smegmatis***

**Supplemental Table 3. RNA oligos**

**Supplemental Table 4. Plasmids used for protein overexpression in BL21 *E. coli***

**Supplemental Table 5. M. smegmatis genes and proteins**

**Supplemental Table 6. Differentially expressed genes with the RNase E IDR deletions**

**Supplemental Table 7. Ribonucleoside reductase genes upregulated in the RNase E double IDR deletion**

## CONFLICT OF INTEREST

None

## FUNDING

This work was supported in part by NIH-NIAID award 5TP01AI143575-02 to SSS, by NIH-NIAID award R21 AI156415-01A1 to SSS, by NSF-CAREER award 1652756 from the Directorate of Biological Sciences to SSS, an award from the Potts Foundation to SSS, a WPI Manning/Leser A&S Faculty Research Grant to SSS, NIH NIGMS award R35GM124733 to JMS, DoEd GAANN training grant P200A240115 to ARR, and a Dr. Armand P. Ferro and Mary H. Ferro Summer Fellowship to ARR. The funders had no role in study design, data collection and analysis, decision to publish, or preparation of the manuscript.

## Notes

### Competing Interest Statement

The authors have declared no competing interest.

### Summary of Updates

RNase E RNA cleavage kinetics experiments were repeated using optimized reaction conditions. The results are consistent with our previous results but are now reported with substantially higher confidence. RNase E RNA binding experiments were repeated using optimized reaction conditions. The results are largely consistent with our previous results except that we now detect RNA binding by the double IDR deletion mutant of RNase E. The rifampicin sensitivity of foci formed by RNase E with various IDR deletions was determined, shedding light on ways that these foci are similar to, and in some cases different from, those formed by full-length RNase E. We repeated several of the coimmunoprecipitation experiments, obtaining more consistent and thus higher-confidence data on the protein-protein interactions that form the mycobacterial degradosome-like network. In addition to what we previously reported, we now conclude that RNase J interacts with both IDRs of RNase E, and that PNPase interacts with the catalytic domain of RNase E as well as its C-terminal IDR. We repeated assays to test for allosteric activation of the helicase RhlE by RNases using optimized conditions and we found that RNase E can stimulate RhlE in a manner that is dependent upon its IDRs. Additional supplemental figures were added and the figures and text revised to improve clarity.

## REFERENCES

1. Organization, W.H. (2025).

2. Kumar, A. and Tenguria, S. (2022) Bacterial survival in the hostile environment. Academic Press Elsevier, Amsterdam, London, England ; San Diego, California.

3. Vargas-Blanco, D.A., Zhou, Y., Zamalloa, L.G., Antonelli, T. and Shell, S.S. (2019) mRNA Degradation Rates Are Coupled to Metabolic Status in Mycobacterium smegmatis. mBio, 10.

4. Sun, H., Vargas-Blanco, D.A., Zhou, Y., Masiello, C.S., Kelly, J.M., Moy, J.K., Korkin, D. and Shell, S.S. (2024) Diverse intrinsic properties shape transcript stability and stabilization in Mycolicibacterium smegmatis. NAR Genom Bioinform, 6, lqae147.

5. Schubert, O.T., Ludwig, C., Kogadeeva, M., Zimmermann, M., Rosenberger, G., Gengenbacher, M., Gillet, L.C., Collins, B.C., Rost, H.L., Kaufmann, S.H. et al. (2015) Absolute Proteome Composition and Dynamics during Dormancy and Resuscitation of Mycobacterium tuberculosis. Cell Host Microbe, 18, 96–108.

6. Zeller, M.E., Csanadi, A., Miczak, A., Rose, T., Bizebard, T. and Kaberdin, V.R. (2007) Quaternary structure and biochemical properties of mycobacterial RNase E/G. Biochem J, 403, 207–215.

7. Taverniti, V., Forti, F., Ghisotti, D. and Putzer, H. (2011) Mycobacterium smegmatis RNase J is a 5’-3’ exo-/endoribonuclease and both RNase J and RNase E are involved in ribosomal RNA maturation. Mol Microbiol, 82, 1260–1276.

8. Unciuleac, M.C., Ghosh, S., de la Cruz, M.J., Goldgur, Y. and Shuman, S. (2021) Structure and mechanism of Mycobacterium smegmatis polynucleotide phosphorylase. RNA, 27, 959–969.

9. Wang, N., Sheng, Y., Liu, Y., Guo, Y., He, J. and Liu, J. (2024) Cryo-EM structures of Mycobacterium tuberculosis polynucleotide phosphorylase suggest a potential mechanism for its RNA substrate degradation. Arch Biochem Biophys, 754, 109917.

10. Zhou, Y., Sun, H., Rapiejko, A.R., Vargas-Blanco, D.A., Martini, M.C., Chase, M.R., Joubran, S.R., Davis, A.B., Dainis, J.P., Kelly, J.M. et al. (2023) Mycobacterial RNase E cleaves with a distinct sequence preference and controls the degradation rates of most Mycolicibacterium smegmatis mRNAs. J Biol Chem, 299, 105312.

11. Plocinski, P., Macios, M., Houghton, J., Niemiec, E., Plocinska, R., Brzostek, A., Slomka, M., Dziadek, J., Young, D. and Dziembowski, A. (2019) Proteomic and transcriptomic experiments reveal an essential role of RNA degradosome complexes in shaping the transcriptome of Mycobacterium tuberculosis. Nucleic Acids Res, 47, 5892–5905.

12. Callaghan, A.J., Marcaida, M.J., Stead, J.A., McDowall, K.J., Scott, W.G. and Luisi, B.F. (2005) Structure of Escherichia coli RNase E catalytic domain and implications for RNA turnover. Nature, 437, 1187–1191.

13. Callaghan, A.J., Redko, Y., Murphy, L.M., Grossmann, J.G., Yates, D., Garman, E., Ilag, L.L., Robinson, C.V., Symmons, M.F., McDowall, K.J. et al. (2005) “Zn-link“: a metal-sharing interface that organizes the quaternary structure and catalytic site of the endoribonuclease, RNase E. Biochemistry, 44, 4667–4675.

14. Moore, C.J., Go, H., Shin, E., Ha, H.J., Song, S., Ha, N.C., Kim, Y.H., Cohen, S.N. and Lee, K. (2021) Substrate-dependent effects of quaternary structure on RNase E activity. Genes Dev, 35, 286–299.

15. Kaberdin, V.R., Miczak, A., Jakobsen, J.S., Lin-Chao, S., McDowall, K.J. and von Gabain, A. (1998) The endoribonucleolytic N-terminal half of Escherichia coli RNase E is evolutionarily conserved in Synechocystis sp. and other bacteria but not the C-terminal half, which is sufficient for degradosome assembly. Proc Natl Acad Sci U S A, 95, 11637–11642.

16. Vanzo, N.F., Li, Y.S., Py, B., Blum, E., Higgins, C.F., Raynal, L.C., Krisch, H.M. and Carpousis, A.J. (1998) Ribonuclease E organizes the protein interactions in the Escherichia coli RNA degradosome. Genes Dev, 12, 2770–2781.

17. Nandana, V., Al-Husini, N., Vaishnav, A., Dilrangi, K.H. and Schrader, J.M. (2024) Caulobacter crescentus RNase E condensation contributes to autoregulation and fitness. Mol Biol Cell, 35, ar104.

18. Cohan, M.C. and Pappu, R.V. (2020) Making the Case for Disordered Proteins and Biomolecular Condensates in Bacteria. Trends Biochem Sci, 45, 668–680.

19. Azaldegui, C.A., Vecchiarelli, A.G. and Biteen, J.S. (2021) The emergence of phase separation as an organizing principle in bacteria. Biophys J, 120, 1123–1138.

20. Collins, M.J., Tomares, D.T., Nandana, V., Schrader, J.M. and Childers, W.S. (2023) RNase E biomolecular condensates stimulate PNPase activity. Sci Rep, 13, 12937.

21. Nandana, V., Rathnayaka-Mudiyanselage, I.W., Muthunayake, N.S., Hatami, A., Mousseau, C.B., Ortiz-Rodriguez, L.A., Vaishnav, J., Collins, M., Gega, A., Mallikaarachchi, K.S. et al. (2023) The BR-body proteome contains a complex network of protein-protein and protein-RNA interactions. Cell Rep, 42, 113229.

22. Al-Husini, N., Tomares, D.T., Pfaffenberger, Z.J., Muthunayake, N.S., Samad, M.A., Zuo, T., Bitar, O., Aretakis, J.R., Bharmal, M.M., Gega, A. et al. (2020) BR-Bodies Provide Selectively Permeable Condensates that Stimulate mRNA Decay and Prevent Release of Decay Intermediates. Mol Cell, 78, 670–682 e678.

23. Hicks, N.D., Yang, J., Zhang, X., Zhao, B., Grad, Y.H., Liu, L., Ou, X., Chang, Z., Xia, H., Zhou, Y. et al. (2018) Clinically prevalent mutations in Mycobacterium tuberculosis alter propionate metabolism and mediate multidrug tolerance. Nat Microbiol, 3, 1032–1042.

24. Martini, M.C., Hicks, N.D., Xiao, J., Alonso, M.N., Barbier, T., Sixsmith, J., Fortune, S.M. and Shell, S.S. (2022) Loss of RNase J leads to multi-drug tolerance and accumulation of highly structured mRNA fragments in Mycobacterium tuberculosis. PLoS Pathog, 18, e1010705.

25. Lengyel, P. (2012) Memories of a senior scientist: on passing the fiftieth anniversary of the beginning of deciphering the genetic code. Annu Rev Microbiol, 66, 27–38.

26. Donovan, W.P. and Kushner, S.R. (1986) Polynucleotide phosphorylase and ribonuclease II are required for cell viability and mRNA turnover in Escherichia coli K-12. Proc Natl Acad Sci U S A, 83, 120–124.

27. Cheng, Z.F., Zuo, Y., Li, Z., Rudd, K.E. and Deutscher, M.P. (1998) The vacB gene required for virulence in Shigella flexneri and Escherichia coli encodes the exoribonuclease RNase R. J Biol Chem, 273, 14077–14080.

28. Zetzsche, H., Raschke, L. and Furtig, B. (2023) Allosteric activation of RhlB by RNase E induces partial duplex opening in substrate RNA. Front Mol Biosci, 10, 1139919.

29. Chandran, V., Poljak, L., Vanzo, N.F., Leroy, A., Miguel, R.N., Fernandez-Recio, J., Parkinson, J., Burns, C., Carpousis, A.J. and Luisi, B.F. (2007) Recognition and cooperation between the ATP-dependent RNA helicase RhlB and ribonuclease RNase E. J Mol Biol, 367, 113–132.

30. Kushner, S.R. (2002) mRNA decay in Escherichia coli comes of age. J Bacteriol, 184, 4658–4665; discussion 4657.

31. Khemici, V. and Carpousis, A.J. (2004) The RNA degradosome and poly(A) polymerase of Escherichia coli are required in vivo for the degradation of small mRNA decay intermediates containing REP-stabilizers. Mol Microbiol, 51, 777–790.

32. Carpousis, A.J., Van Houwe, G., Ehretsmann, C. and Krisch, H.M. (1994) Copurification of E. coli RNAase E and PNPase: evidence for a specific association between two enzymes important in RNA processing and degradation. Cell, 76, 889–900.

33. Py, B., Causton, H., Mudd, E.A. and Higgins, C.F. (1994) A protein complex mediating mRNA degradation in Escherichia coli. Mol Microbiol, 14, 717–729.

34. Py, B., Higgins, C.F., Krisch, H.M. and Carpousis, A.J. (1996) A DEAD-box RNA helicase in the Escherichia coli RNA degradosome. Nature, 381, 169–172.

35. Miczak, A., Kaberdin, V.R., Wei, C.L. and Lin-Chao, S. (1996) Proteins associated with RNase E in a multicomponent ribonucleolytic complex. Proc Natl Acad Sci U S A, 93, 3865–3869.

36. Worrall, J.A., Gorna, M., Crump, N.T., Phillips, L.G., Tuck, A.C., Price, A.J., Bavro, V.N. and Luisi, B.F. (2008) Reconstitution and analysis of the multienzyme Escherichia coli RNA degradosome. J Mol Biol, 382, 870–883.

37. Prud’homme-Genereux, A., Beran, R.K., Iost, I., Ramey, C.S., Mackie, G.A. and Simons, R.W. (2004) Physical and functional interactions among RNase E, polynucleotide phosphorylase and the cold-shock protein, CsdA: evidence for a ‘cold shock degradosome’. Mol Microbiol, 54, 1409–1421.

38. Lopez, P.J., Marchand, I., Joyce, S.A. and Dreyfus, M. (1999) The C-terminal half of RNase E, which organizes the Escherichia coli degradosome, participates in mRNA degradation but not rRNA processing in vivo. Mol Microbiol, 33, 188–199.

39. Khemici, V., Poljak, L., Luisi, B.F. and Carpousis, A.J. (2008) The RNase E of Escherichia coli is a membrane-binding protein. Mol Microbiol, 70, 799–813.

40. Hardwick, S.W., Chan, V.S., Broadhurst, R.W. and Luisi, B.F. (2011) An RNA degradosome assembly in Caulobacter crescentus. Nucleic Acids Res, 39, 1449–1459.

41. Aguirre, A.A., Vicente, A.M., Hardwick, S.W., Alvelos, D.M., Mazzon, R.R., Luisi, B.F. and Marques, M.V. (2017) Association of the Cold Shock DEAD-Box RNA Helicase RhlE to the RNA Degradosome in Caulobacter crescentus. J Bacteriol, 199.

42. Commichau, F.M., Rothe, F.M., Herzberg, C., Wagner, E., Hellwig, D., Lehnik-Habrink, M., Hammer, E., Volker, U. and Stulke, J. (2009) Novel activities of glycolytic enzymes in Bacillus subtilis: interactions with essential proteins involved in mRNA processing. Mol Cell Proteomics, 8, 1350–1360.

43. Lehnik-Habrink, M., Newman, J., Rothe, F.M., Solovyova, A.S., Rodrigues, C., Herzberg, C., Commichau, F.M., Lewis, R.J. and Stulke, J. (2011) RNase Y in Bacillus subtilis: a Natively disordered protein that is the functional equivalent of RNase E from Escherichia coli. J Bacteriol, 193, 5431–5441.

44. Cascante-Estepa, N., Gunka, K. and Stulke, J. (2016) Localization of Components of the RNA-Degrading Machine in Bacillus subtilis. Front Microbiol, 7, 1492.

45. Durand, S. and Condon, C. (2018) RNases and Helicases in Gram-Positive Bacteria. Microbiol Spectr, 6.

46. Oviedo-Bocanegra, L.M., Hinrichs, R., Rotter, D.A.O., Dersch, S. and Graumann, P.L. (2021) Single molecule/particle tracking analysis program SMTracker 2.0 reveals different dynamics of proteins within the RNA degradosome complex in Bacillus subtilis. Nucleic Acids Res, 49, e112.

47. Tejada-Arranz, A., Galtier, E., El Mortaji, L., Turlin, E., Ershov, D. and De Reuse, H. (2020) The RNase J-Based RNA Degradosome Is Compartmentalized in the Gastric Pathogen Helicobacter pylori. mBio, 11.

48. Ehrt, S., Guo, X.V., Hickey, C.M., Ryou, M., Monteleone, M., Riley, L.W. and Schnappinger, D. (2005) Controlling gene expression in mycobacteria with anhydrotetracycline and Tet repressor. Nucleic Acids Res, 33, e21.

49. Culviner, P.H., Guegler, C.K. and Laub, M.T. (2020) A Simple, Cost-Effective, and Robust Method for rRNA Depletion in RNA-Sequencing Studies. mBio, 11.

50. Culviner, P.H. and Laub, M.T. (2018) Global Analysis of the E. coli Toxin MazF Reveals Widespread Cleavage of mRNA and the Inhibition of rRNA Maturation and Ribosome Biogenesis. Mol Cell, 70, 868–880 e810.

51. Li, H. and Durbin, R. (2009) Fast and accurate short read alignment with Burrows-Wheeler transform. Bioinformatics, 25, 1754–1760.

52. Love, M.I., Huber, W. and Anders, S. (2014) Moderated estimation of fold change and dispersion for RNA-seq data with DESeq2. Genome Biol, 15, 550.

53. Hulsen, T., de Vlieg, J. and Alkema, W. (2008) BioVenn - a web application for the comparison and visualization of biological lists using area-proportional Venn diagrams. BMC Genomics, 9, 488.

54. Rapiejko, A.R., Reddy, M., Sacchettini, J.C. and Shell, S.S. (2026) In vitro cleavage requirements and specificities of mycobacterial RNase E. Biochem Biophys Res Commun, 828, 154079.

55. Ait-Bara, S. and Carpousis, A.J. (2015) RNA degradosomes in bacteria and chloroplasts: classification, distribution and evolution of RNase E homologs. Mol Microbiol, 97, 1021–1135.

56. Krishnamoorthy, G., Kaiser, P., Lozza, L., Hahnke, K., Mollenkopf, H.J. and Kaufmann, S.H.E. (2019) Mycofactocin Is Associated with Ethanol Metabolism in Mycobacteria. mBio, 10.

57. Jain, P., Malakar, B., Khan, M.Z., Lochab, S., Singh, A. and Nandicoori, V.K. (2018) Delineating FtsQ-mediated regulation of cell division in Mycobacterium tuberculosis. J Biol Chem, 293, 12331–12349.

58. Pickford, H., Alcock, E., Singh, A., Kelemen, G. and Bhatt, A. (2020) A mycobacterial DivIVA domain-containing protein involved in cell length and septation. Microbiology (Reading), 166, 817–825.

59. Wu, K.J., Zhang, J., Baranowski, C., Leung, V., Rego, E.H., Morita, Y.S., Rubin, E.J. and Boutte, C.C. (2018) Characterization of Conserved and Novel Septal Factors in Mycobacterium smegmatis. J Bacteriol, 200.

60. Bandyra, K.J., Wandzik, J.M. and Luisi, B.F. (2018) Substrate Recognition and Autoinhibition in the Central Ribonuclease RNase E. Mol Cell, 72, 275–285 e274.

61. J. Hilario Cafiero, J.X., Irene Lepori, Abigail R. Rapiejko, Manchi Reddy, Louis A. Roberts, M. Sloan Siegrist, James C. Sacchettini, Scarlet S. Shell. (2025) ycobacterium tuberculosis MutT4 is an RNA pyrophosphohydrolase that forms biomolecular condensates and sensitizes mRNAs to degradation. bioRxiv.

62. Burck, M., Nouaille, S., Carpousis, A.J., Cocaign-Bousquet, M. and Girbal, L. (2025) rRNA loss induced by rifampicin addition in Escherichia coli reflects extraction artifacts rather than in vivo degradation. Sci Rep, 15, 30833.

63. Griego, A., Douche, T., Gianetto, Q.G., Matondo, M. and Manina, G. (2022) RNase E and HupB dynamics foster mycobacterial cell homeostasis and fitness. iScience, 25, 104233.

64. Spoelstra, W.K., van der Sluis, E.O., Dogterom, M. and Reese, L. (2020) Nonspherical Coacervate Shapes in an Enzyme-Driven Active System. Langmuir, 36, 1956–1964.

65. Hausmann, S., Gonzalez, D., Geiser, J. and Valentini, M. (2021) The DEAD-box RNA helicase RhlE2 is a global regulator of Pseudomonas aeruginosa lifestyle and pathogenesis. Nucleic Acids Res, 49, 6925–6940.

66. Rock, J.M., Hopkins, F.F., Chavez, A., Diallo, M., Chase, M.R., Gerrick, E.R., Pritchard, J.R., Church, G.M., Rubin, E.J., Sassetti, C.M. et al. (2017) Programmable transcriptional repression in mycobacteria using an orthogonal CRISPR interference platform. Nat Microbiol, 2, 16274.

67. Belton, M., Brilha, S., Manavaki, R., Mauri, F., Nijran, K., Hong, Y.T., Patel, N.H., Dembek, M., Tezera, L., Green, J. et al. (2016) Hypoxia and tissue destruction in pulmonary TB. Thorax, 71, 1145–1153.

68. Via, L.E., Lin, P.L., Ray, S.M., Carrillo, J., Allen, S.S., Eum, S.Y., Taylor, K., Klein, E., Manjunatha, U., Gonzales, J. et al. (2008) Tuberculous granulomas are hypoxic in guinea pigs, rabbits, and nonhuman primates. Infect Immun, 76, 2333–2340.

69. Ono, M. and Kuwano, M. (1979) A conditional lethal mutation in an Escherichia coli strain with a longer chemical lifetime of messenger RNA. J Mol Biol, 129, 343–357.

70. Clarke, J.E., Kime, L., Romero, A.D. and McDowall, K.J. (2014) Direct entry by RNase E is a major pathway for the degradation and processing of RNA in Escherichia coli. Nucleic Acids Res, 42, 11733–11751.

71. Hammarlof, D.L., Bergman, J.M., Garmendia, E. and Hughes, D. (2015) Turnover of mRNAs is one of the essential functions of RNase E. Mol Microbiol, 98, 34–45.

72. Santonoceto, G., Jurkiewicz, A. and Szczesny, R.J. (2024) RNA degradation in human mitochondria: the journey is not finished. Hum Mol Genet, 33, R26–R33.

73. Teixeira, D. and Parker, R. (2007) Analysis of P-body assembly in Saccharomyces cerevisiae. Mol Biol Cell, 18, 2274–2287.

74. Currie, S.L., Xing, W., Muhlrad, D., Decker, C.J., Parker, R. and Rosen, M.K. (2023) Quantitative reconstitution of yeast RNA processing bodies. Proc Natl Acad Sci U S A, 120, e2214064120.

