## Supplemental Figures for "Defining the role of RNase E in the mycobacterial degradosome-like network"

```
eco            ------------------------------------------------------------   0
abscessus      MADYELPEDLHTSAPEGEHAALTPDTPPAPAEQGELPEKLRVHSLARVLGTSTRRVLDAL   60
mtb            ----------------MIDGAP-PSDPPEPSQHEELPDRLRVHSLARTLGTTSRRVLDAL   43
smeg           ----------------MAEDAHTEDLSTQTPQQEGLPERLRVHSLARVLGTTSRRVLDAL   44

eco            ------------------------------------------------------------   0
abscessus      SNLDGRARSPHSNVDKQDAAKVREVLANEAEAAPVVAETPAPVAAEEA--------PAVV   112
mtb            TALDGRVRSAHSTVDRVDAVRVRDLLATHLET------------------------AGVL   79
smeg           AEFDGRQRSAHSTVDKADAERVRAALTESPAAETPPEEAPAAETPVADLVVVQAEQVEVV   104

eco            ------------------------------------------------------------   0
abscessus      TGD---------------------------SADSAEDPDNPPTTPILI----AEATVIEA   141
mtb            AASVH---------------------------------------APEASEEPESRLMLET   100
smeg           TVSEAGPAEPAEPAEPEAPAAEAEAEAETEVADEAETPEPTFRGAVLVGDEPESRLILEH   164

eco            ------------------------------------------------------------   0
abscessus      AAVPVPAAAAPTAYDYGPLFVAPQAPEPQERPAPVRTEKAEDVDEDAGE-----TDDSDA   196
mtb            Q---ETRNADVERPHYMPLFVAPQPIPE---PLA--DDEDVDDGP------DYVADDSDA   146
smeg           ANIPPARETQTERPDYLPLFVAPQPVSF---EPAVVDDEDEDDDTETGAESDFDSGADSD   221

eco            ------------------------------------------------------------   0
abscessus      DGEGDGDRP-ARRRRRGRRGRGRGRGEQGGGEGSDDSGDDAAAE-GGEAAEGDDAADGTG   254
mtb            DDEGQLDRPANRRRRRGRRGRGRGRGEQGGSDGDPVDQQSEPRAQQFTSADAAETDDGDD   206
smeg           SDDDQADR--PRRRRRGRRGRGRGRGEQNDDATSDADTDSTE-----------DQTDGDE   268

eco            ------------------------------------------------------------   0
abscessus      DDSPAPDGDESNAEDGDTPEGASRRRRRRRRRKAGSGDDGDG----PSDDDPPNTVVHER   310
mtb            RDSEDTEAGDNGEDENGSLEAGNRRRRRRRRRKSASGDDNDAALEGPLPDDPPNTVVHER   266
smeg           QE-SGEDSDDSGDEDSTTTEGGTRRRRRRRRRKSGSGDSDDA----VSPDDPPNTVVHER   323

eco            ------------------------------------------------------------   0
abscessus      KPRAERG-----------DEIQGISGSTRLEAKRQRRRDGRDAGRRRPPILTEAEFLARR   359
mtb            VPRAGDKAGNSQDGGSGSTEIKGIDGSTRLEAKRQRRRDGRDAGRRRPPVLSEAEFLARR   326
smeg           APRTERS------DKSDDSEIQGISGSTRLEAKRQRRRDGRDAGRRRPPILSEAEFLARR   377
                     ↑

eco            --MKR-MLINATQ--------QEELRVALVDGQRLYDLDIESPGHEQKKANIYKGKITRI   49
abscessus      EAVDRVMVVRDKTRSEPPHEGARYTQIAVLEDGVLVEHFVTSAASASLVGNIYLGVVQNV   419
mtb            EAVERVMVVRDRVRTEPPLPGTRYTQIAVLEDGIVVEHFVTSAASASLVGNIYLGIVQNV   386
smeg           EAVERTMIVRDKVRTEPPHEGARYTQIAVLEDGVVVEHFVTSAASASLVGNIYLGIVQNV   437
                 :.* *::.            .  ::*:::.  : :  : * .  .  .*** * : .:

eco            EPSLEAAFVDYGAERHGFLPLKEIAREYFPANYSAHGRPNIKDVLREGQEVIVQIDKEER   109
abscessus      LPSMEAAFVDIGRGRNGVLYAGEVNWDA---AGLGGSNRKIEQALKSGDYVLVQVSKDPV   476
mtb            LPSMEAAFVDIGRGRNGVLYAGEVNWDA---AGLGGADRKIEQALKPGDYVVVQVSKDPV   443
smeg           LPSMEAAFVDIGRGRNGVLYAGEVNWEA---AGLGGQNRKIEQALKPGDYVVVQVSKDPV   494
                **:****** *  *:*.*   *:  :       .    :*::.*: *: *:**:.*:
```

**Supplementary Figure 1 (continued)**

```
eco          GNKGAALTTFISLAGSYLVLMPNNPRAGGISRRIEGDDRTELKEALASLELPEGMGLIVR   169
abscessus    GHKGARLTTQISLAGRYLVYVPGAS-STGISRKLPDTERQRLKEILRDI-VPADAGVIIR   534
mtb          GHKGARLTTQVSLAGRFLVYVPGAS-STGISRKLPDTERQRLKEILREV-VPSDAGVIIR   501
smeg         GHKGARLTTQVSLAGRYLVYVPGAS-STGISRKLPDTERQRLKEILREV-VPSDAGVIIR   552
             *:*** *** :**** :** :*.   : ****:: . :* .*** * .: :* . *:*:*

eco          TAGVGKSAEALQWDLSFRLKHWEAIKKAAES-----RPAPFLIHQESNVIVRAFRDYLRQ   224
abscessus    TASEGVREEDIRADVQRLQEQWQGIEQSATDLTSKASGAAVALYEEPDVLVKVVRDLFNE   594
mtb          TASEGVKEDDIRADVARLRERWEQIEAKAQETKEKAAGAAVALYEEPDVLVKVIRDLFNE   561
smeg         TASEGVKEEDIRSDVERLQKRWSEIEAKAAEVTEKKAGAAVALYEEPDVLVKVIRDLFNE   612
             **. *   : :: *:    ::*. *:  * .       * . :::* :*:*:..** :.:

eco          DIGEILIDNPKVLELARQHIAALGRPDFSSKIKLYTG--------EIPLFSHYQIESQIE   276
abscessus    DFSKLIIEGDAAWDTIDDYVKT-VAPDLLSRIERYESGEG------PDVFAVHRIDEQLA   647
mtb          DFVGLIVSGDEAWNTINEYVNS-VAPELVSKLTKYESADGPDGQSAPDVFTVHRIDEQLA   620
smeg         DFSSLIVSGDEAWNTINSYVEA-VAPDLMPRLTKYEP----AGPDAPDVFAVHRIDEQLA   667
             *:  :::..  . :   .:: :   *::  ::  *             :*: ::*:.*:

eco          SAFQREVRLPSGGSIVIDSTEALTAIDINSARAT-RGGDIEETAFNTNLEAADEIARQLR   335
abscessus    KALDRKVWLPSGGTLVIDRTEAMTVVDVNTGKFTGAGGNLEETVTRNNLEAAEEIVRQLR   707
mtb          KAMDRKVWLPSGGTLVIDRTEAMTVIDVNTGKFTGAGGNLEQTVTKNNLEAAEEIVRQLR   680
smeg         KAMDRKVWLPSGGTLVIDRTEAMTVVDVNTGKFTGSGGNLEQTVTRNNLEAAEEIVRQLR   727
             .*::*:* *****::*** ***:*.:*:*:.: *  **::*:*. ..*****:**.****

eco          LRDLGGLIVIDFIDMTPVRHQRAVENRLREAVRQDRARIQISHISRFGLLEMSRQRLSPS   395
abscessus    LRDVGGIVVIDFIDMVLESNRDLVLRRLTEALGRDRTRHQVSEVTSLGLVQLTRKKLGTG   767
mtb          LRDIGGIVVIDFIDMVLESNRDLVLRRLTESLARDRTRHQVSEVTSLGLVQLTRKRLGTG   740
smeg         LRDIGGIVVIDFIDMVLESNRDLVLRRLTEALARDRTRHQVSEVTSLGLVQLTRKRLGTG   787
             ***:**::*******.   ::  * .** *:: :**:* *:*.:: :**::::*::*. .

eco          LGESSHHVCPRCSGTGTVRDNESLSLSILRLIEEEALKENTQEVHAIVPVPIASYLLNEK   455
abscessus    LIEAFSSTCTHCAGRGIVLHVDPVDNASNASSPAKK-------------AE---TGRRAK   811
mtb          LIEAFSTSCPNCSGRGILLHADPVDSAAATGRKSE---------------P---GARRGK   782
smeg         LVEAFSTACTHCGGRGIVLHGDPIDSASSNGGRKSDSSGGG-------GSG---GGRRGK   837
                                                    ↑
             * *:    * .*.* * : . : :. :        .                   . *

eco          RSAVNAIETRQDGVRCVIVPN-DQMETPHYHVLRVRKGEETPTLSYMLPKLHEEAMALPS   514
abscessus    RSRKG----KTDDVVVARTPSHPQGEHPMFKAMAAANGHHEDDEDA---DVNEDAL----   860
mtb          RSKKSRSEESSDRSMVAKVPVHAPGEHPMFKAMAAGLSSLAGRGDE---ESGEPAA----   835
smeg         RGKKG--AARTEEVHVAKVPDHTPGEHPMFKAMAAANGKHEGDEDH---EDHEDHE----   888
             *.  .      :    . .*    * * ::.: .  .       .    .  *

eco          EEEFAERKRPEQPALATFAMPDVPPAPTPAEPAAPVVAPAPKAAPATPAAPAQPGLLSRF   574
abscessus    -----D-----DDALEEVE-----------EVEEIVETIQSAET--------PDNVP--   888
mtb          -----E-------------LAEQ----------------AGDQP--------PTDLDDT   852
smeg         -----T-----AEDTTAAEVRDDTRDEHDADERAHVVTAAVGAAG--------DEDLDDS   930
                                                                    :
```

**Supplementary Figure 1 (continued)**

```
eco             FGALKALFSGGEETKPTEQPAPKAEAKPERQQDRRKPRQNNRRDRNERRDTRSERTEGSD    634
abscessus       ----------------DQ----------------DEPE--HEPA----SAGSN-------    903
mtb             A---QADFEDTEDTDEDED---ELDADE-DLEDLDDEDLDEDLD----VEDS-------D    894
smeg            DEDSDLDSDEESDDESDED---EIELDD------DEDELDEDIE----VIGDSDDSDDSD    977
                                 :                 .    .

eco             NREENRR-NRRQAQQQTAETRESRQQAEVTEKARTADEQQAPRRERSRRRNDDKRQAQQE    693
abscessus       AASEP-----WAEGAPEAAATEP-------------G-TAEPEA----------------    928
mtb             SDDEDSDEDAADADVDEEDA--A-------------GLDGSP-G----------------    922
smeg            DSDEDDDSDDSDDDSDEDEDSDS-------------DEDEEPVR----------------    1008
                  .*                                .    *

eco             AKALNVEEQSVQETEQEERVRPVQPRRKQRQLNQKVRYEQSVAEEAVVAPVVEETVAAEP    753
abscessus       -------PAATPEPAAAAPTRPRRRRSAARAAGPPVEH----------------------    959
mtb             -------EVDVPGVTELAPTR-PRRRVAGRPAGPPIRLD---------------------    953
smeg            -------EVYEPPV-TAPRAR-VRRRAAARPAGPPSHD----------------------    1037
                                   .*  : *   *  .   .

eco             IVQEAPAPRTELVKVPLPVVAQTAPEQQEENNADNRDNGGMPRRSRRSPRHLRVSGQRRR    813

eco             RYRDERYPTQSPMPLTVACASPELASGKVWIRYPIVRPQDVQVEEQREQEEVHVQPMVTE    873

eco             VPVAAAIEPVVSAPVVEEVAGVVEAPVQVAEPQPEVVETTHPEVIAAAVTEQPQVITESD    933

eco             VAVAQEVAEQAEPVVEPQEETADIEEVVETAEVVVAEPEVVAQPAAPVVAEVAAEVETVA    993

eco             AVEPEVTVEHNHATAPMTRAPAPEYVPEAPRHSDWQRPTFAFEGKGAAGGHTATHHASAA    1053

eco             PARPQPVE 1061
```

**Supplementary Figure 1. Clustal Omega multiple sequence alignment of RNase E from *E. coli* (eco), *M. abscessus* (abscessus), *M. tuberculosis* (Mtb), and *M. smegmatis* (smeg).**

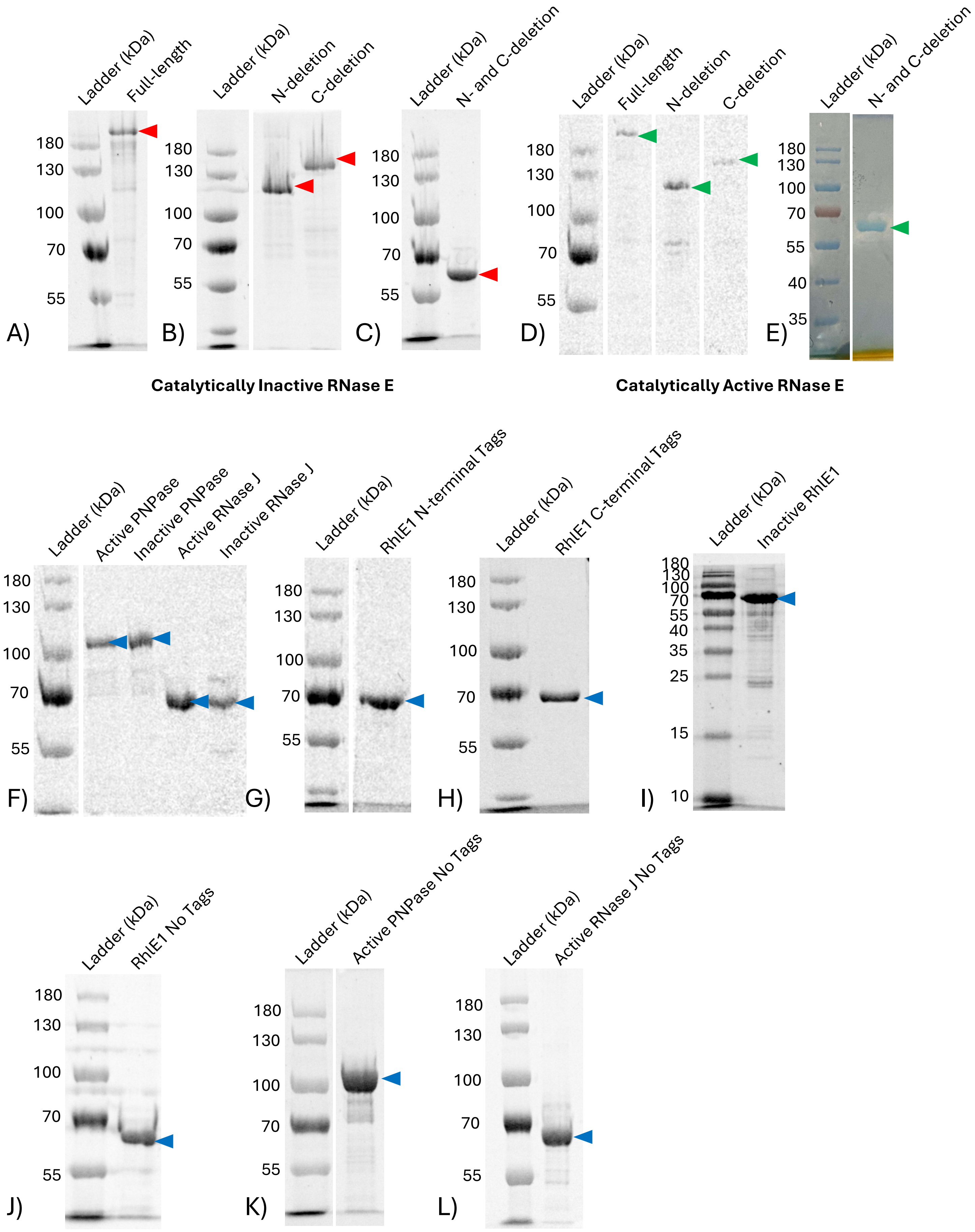
Ladder (kDa)
Full-length
180
130
100
70
55
A)
Ladder (kDa)
N-deletion
C-deletion
180
130
100
70
55
B)
Ladder (kDa)
N- and C-deletion
180
130
100
70
55
C)
Ladder (kDa)
Full-length
N-deletion
C-deletion
180
130
100
70
55
D)
Ladder (kDa)
N- and C-deletion
180
130
100
70
55
40
35
E)
Catalytically Inactive RNase E
Catalytically Active RNase E
Ladder (kDa)
Active PNPase
Inactive PNPase
Active RNase J
Inactive RNase J
180
130
100
70
55
F)
Ladder (kDa)
RhlE1 N-terminal Tags
180
130
100
70
55
G)
Ladder (kDa)
RhlE1 C-terminal Tags
180
130
100
70
55
H)
Ladder (kDa)
Inactive RhlE1
180
130
100
70
55
40
35
25
15
10
I)
Ladder (kDa)
RhlE1 No Tags
180
130
100
70
55
J)
Ladder (kDa)
Active PNPase No Tags
180
130
100
70
55
K)
Ladder (kDa)
Active RNase J No Tags
180
130
100
70
55
L)

**Supplementary Figure 2.** Representative preparations of purified proteins. 3000 ng of total purified protein was run on a 7.5% or 10% SDS-PAGE gels, as appropriate. Arrows indicate the band of the protein of interest. **A)** Full-length, **B)** N-deletion and C-deletion, and **C)** N- and C-double deletion RNase E are shown with the catalytically inactive version. **D)** Full-length, N-deletion, and C-deletion and **E)** N-and C-double deletion RNase E are shown with the catalytically active version. **F)** shows catalytically active PNPase and RNase J and catalytically inactive PNPase and RNase J with HA and his tags. **G)** shows RhlE1 with N-terminal HA and his tags. **H)** shows RhlE1 with C-terminal HA and his tags. **I)** shows inactive RhlE. **J)** shows RhlE, **K)** shows active PNPase, and **L)** shows active RNase J with his tag only (no HA tag).

A)

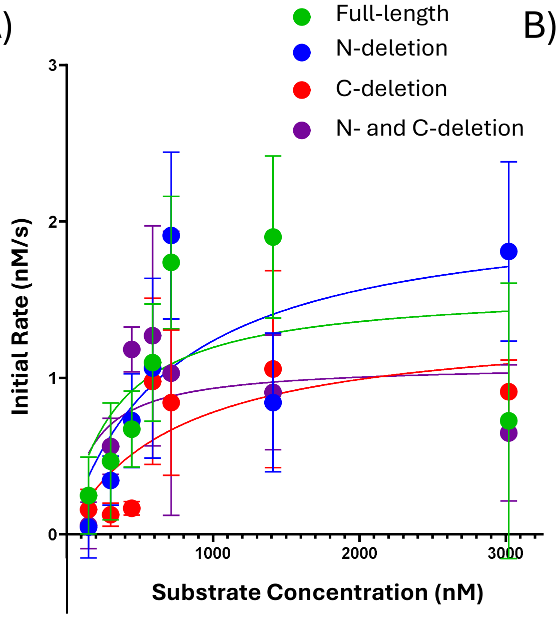


B)

| | Apparent $K_m$ (nM) | $K_m$ 95% CI | Apparent $V_{max}$ (nM/s) | Apparent $K_{cat}$ ($s^{-1}$) | $K_{cat}$ 95% CI | P-value |
|---|---|---|---|---|---|---|
| **Full-length** | 322 | 33 to 1144 | 1.58 | 0.016 | 0.0095 to 0.025 | - |
| **N-deletion** | 710. | 176 to 2940 | 2.12 | 0.021 | 0.013 to 0.042 | 0.658 |
| **C-deletion** | 737 | 161 to 3050 | 1.35 | 0.013 | 0.0077 to 0.027 | 0.0780 |
| **N- and C-deletion** | 166 | -15 to 710 | 1.09 | 0.011 | 0.0066 to 0.017 | 0.447 |

**Supplemental Figure 3. A)** Preliminary kinetic analysis of IDR deletion RNase E mutants *in vitro* using a 29 nt FAM labeled substrate, quantified in **B).** Initial rates were calculated by simple linear regression of the disappearance of substrate, quantified on ImageJ with the FIJI plugin. Points represent the average of three replicates. Nonlinear regression with a $K_{cat}$ model was used for determining kinetic constants, and each mutant was compared to full-length using the extra sum of squares F-test.

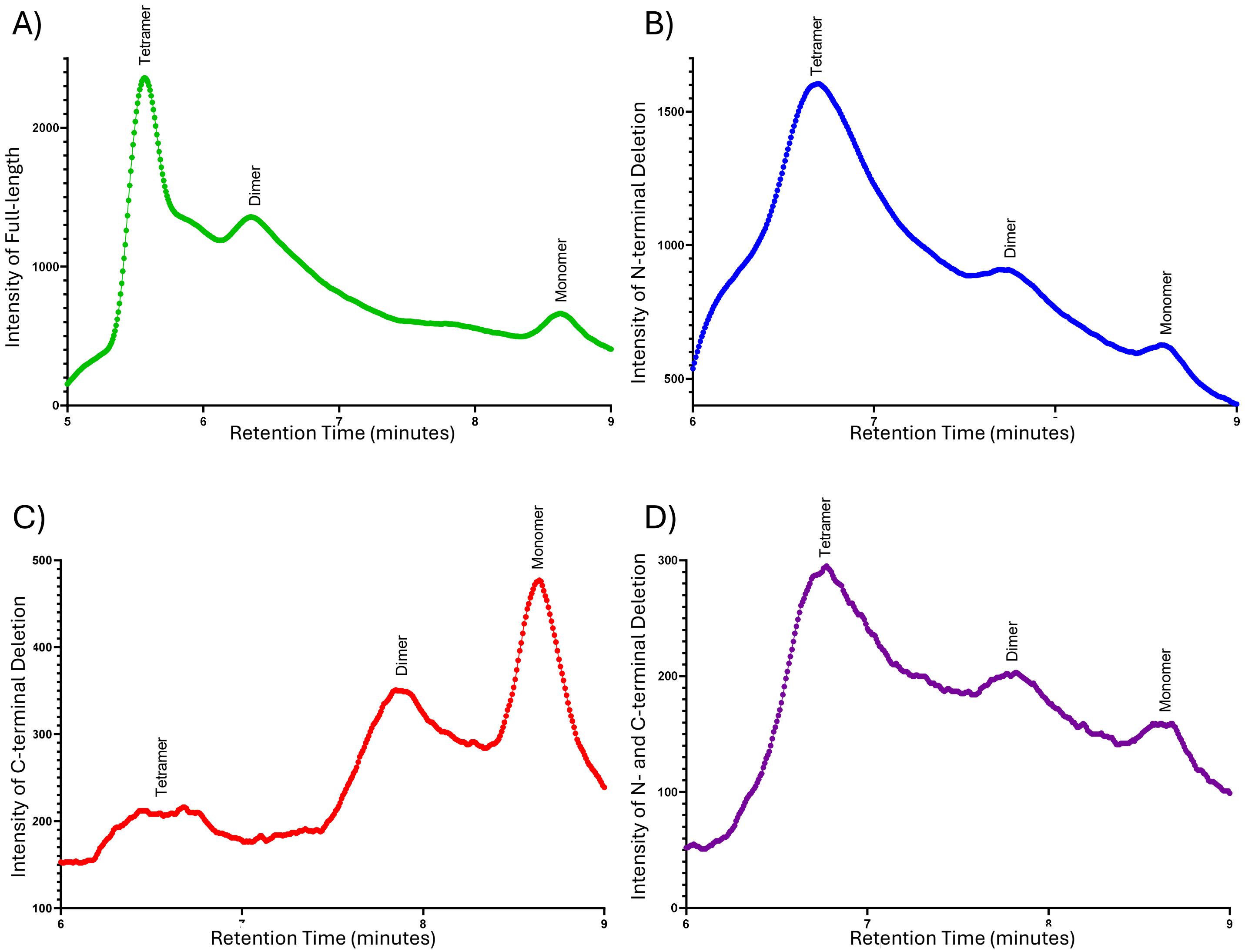


**Supplementary Figure 4.** Representative chromatograms of analytical size exclusion chromatography on a high-performance liquid chromatography system for catalytically inactive **A)** full-length, **B)** N-deletion, **C)** C-deletion, and **D)** N- and C-deletion RNase E. A molecular weight standard was used to estimate the relative size of the peaks based on retention time to determine multimerization status.

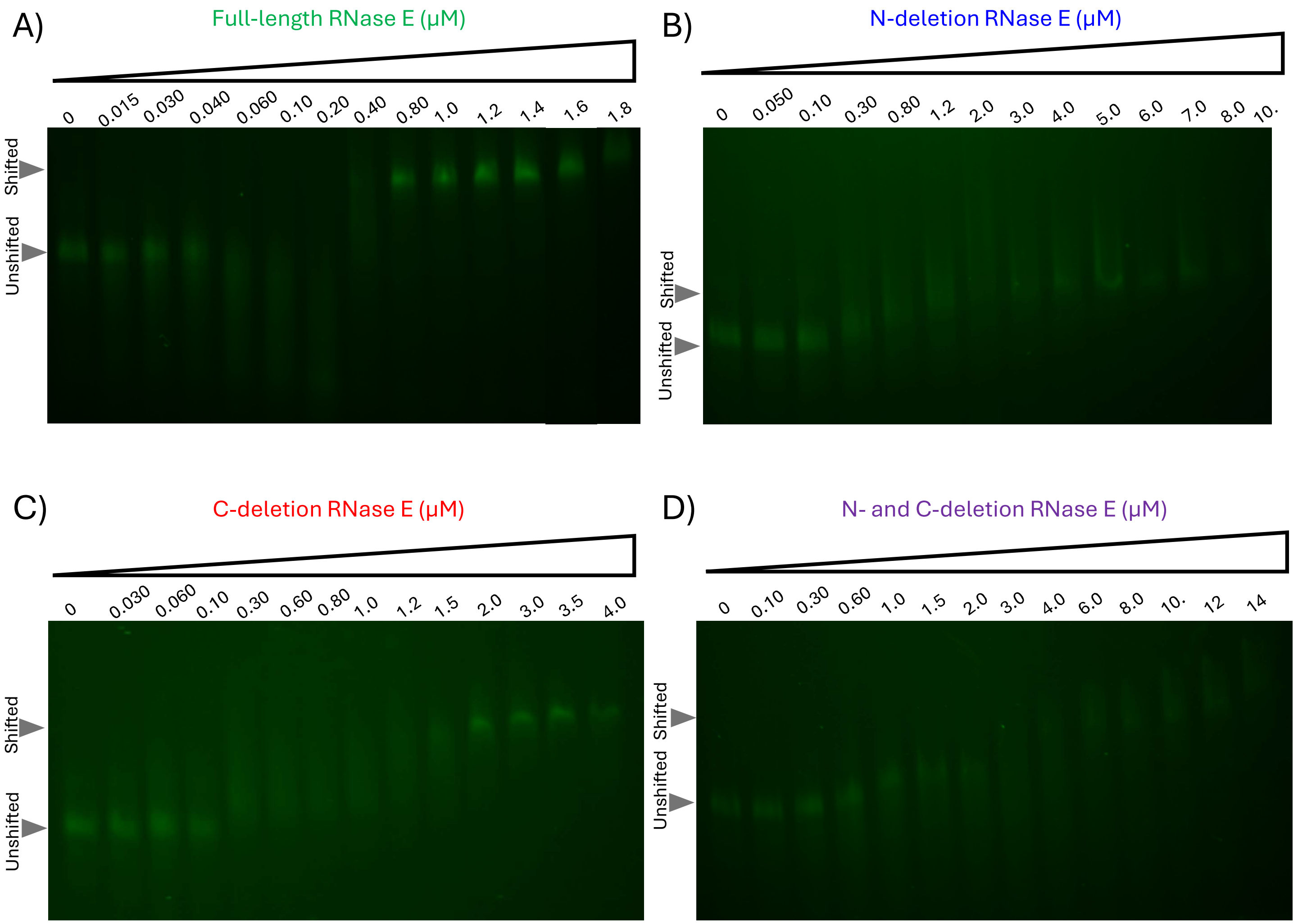


**Supplemental Figure 5.** Representative EMSA native gels used for quantification of shift of a 721 nt fluorescein-labeled RNA with **A)** full-length, **B)** N-deletion, **C)** C-deletion, and **D)** N- and C-deletion catalytically inactive RNase E.

A)

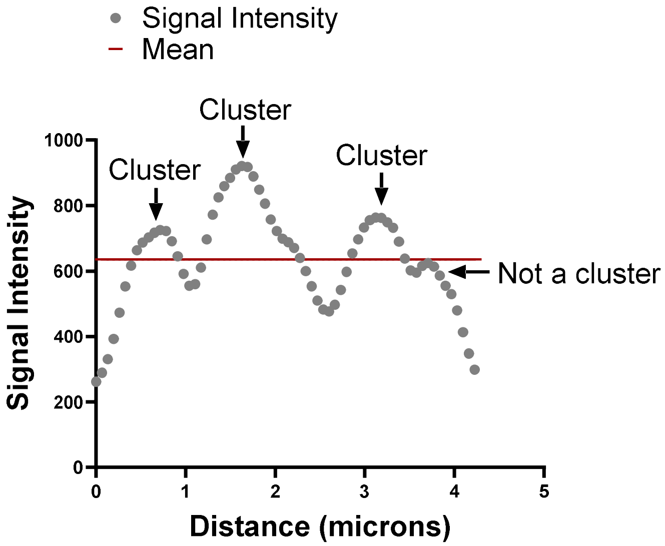


**Supplementary Figure 6.** Example to demonstrate microscopy quantification methods. A line was drawn through the long axis of each cell in FIJI, and the signal intensity was obtained. **A)** shows the quantification method for calculating clusters per micron of fluorescently tagged proteins. Clusters were considered areas where the signal rose above then dipped below the mean, as indicated. Areas that did rise above the mean (as shown) were not considered clusters by this method. The number of clusters was normalized by cell length and reported in Figure 4B, E, and H and Figure 7A.

A)

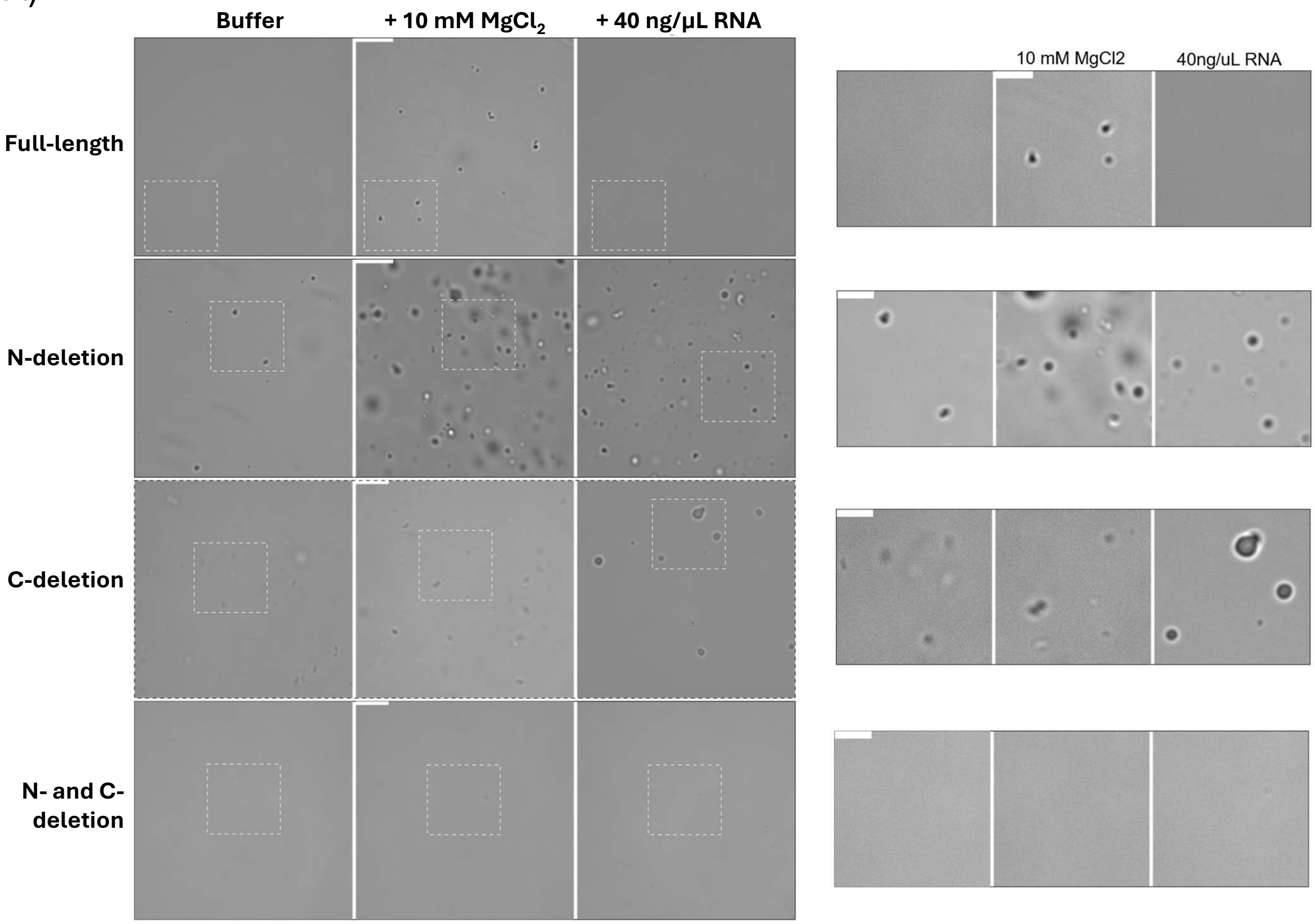


B)

| | Buffer | + 10 mM $MgCl_2$ | + RNA |
|---|---|---|---|
| Full-length | - | + | - |
| N-deletion | + | + | + |
| C-deletion | + | + | + |
| N- and C- deletion | - | - | - |

C)

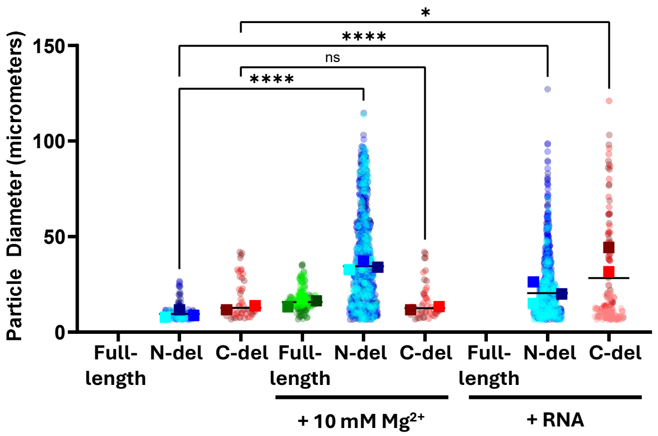


**Supplementary Figure 7. A)** Representative microscopy images of *in vitro* condensate-like body formation by RNase E and its IDR deletions in buffer composed of 20 mM Tris pH 7.4, 100 mM NaCl, 1 mM DTT, and 1 mM $Mg^{2+}$, with the $Mg^{2+}$ concentration increased to 10 mM or total *E. coli* RNA added where indicated. Catalytically active proteins were used in these experiments. Scalebars represent 19.5 µm (main panels, left) or 9.75 µm (magnified regions, right). **B)** Chart indicating which proteins and conditions showed *in vitro* condensate formation. **C)** Superplot showing the pixel diameter of *in vitro* condensates in each condition. The circle colors represent biological replicates, and the squares represent the median of that biological replicate. Black lines represents the means of the medians. Pixel diameter could not be measured for conditions in which condensates did not form. A one-way ANOVA with Sidak's multiple comparisons test was performed. ns $p > 0.05$, * $p \leq 0.05$, **** $p \leq 0.0001$

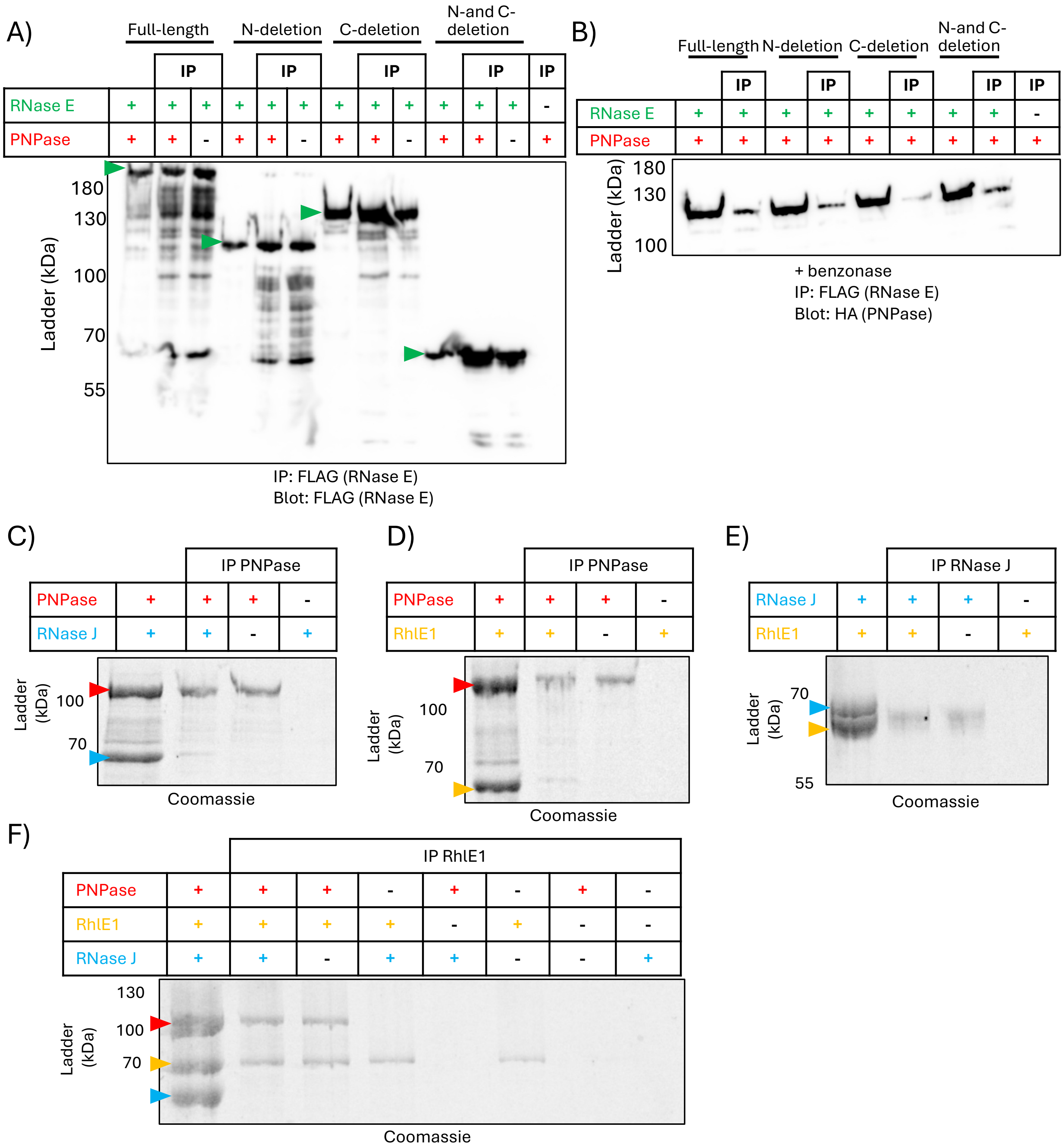


**Supplementary Figure 8. A)** Representative image of an anti-FLAG western blot detecting FLAG-tagged RNase E in an anti-FLAG pulldown. **B)** Representative image of four independent replicates of a pulldown of FLAG-tagged RNase E and coimmunoprecipitation of HA-tagged PNPase with benzonase to remove any RNA bridges, detected by western blot. In all cases "IP" indicates immunoprecipitation, and samples incubated in buffer without beads were run in parallel for comparison. Representative images of Coomassie stained SDS-PAGE gels of the HA pull downs with **C)** HA-PNPase and untagged RNase J, **D)** HA-PNPase and untagged RhlE1, **E)** HA-RNase J and untagged RhlE1, and **F)** a three-protein HA coimmunoprecipitation between HA-RhlE and untagged PNPase and untagged RNase J. In all cases "IP" indicates immunoprecipitation, and samples incubated in buffer without beads were run in parallel for comparison.

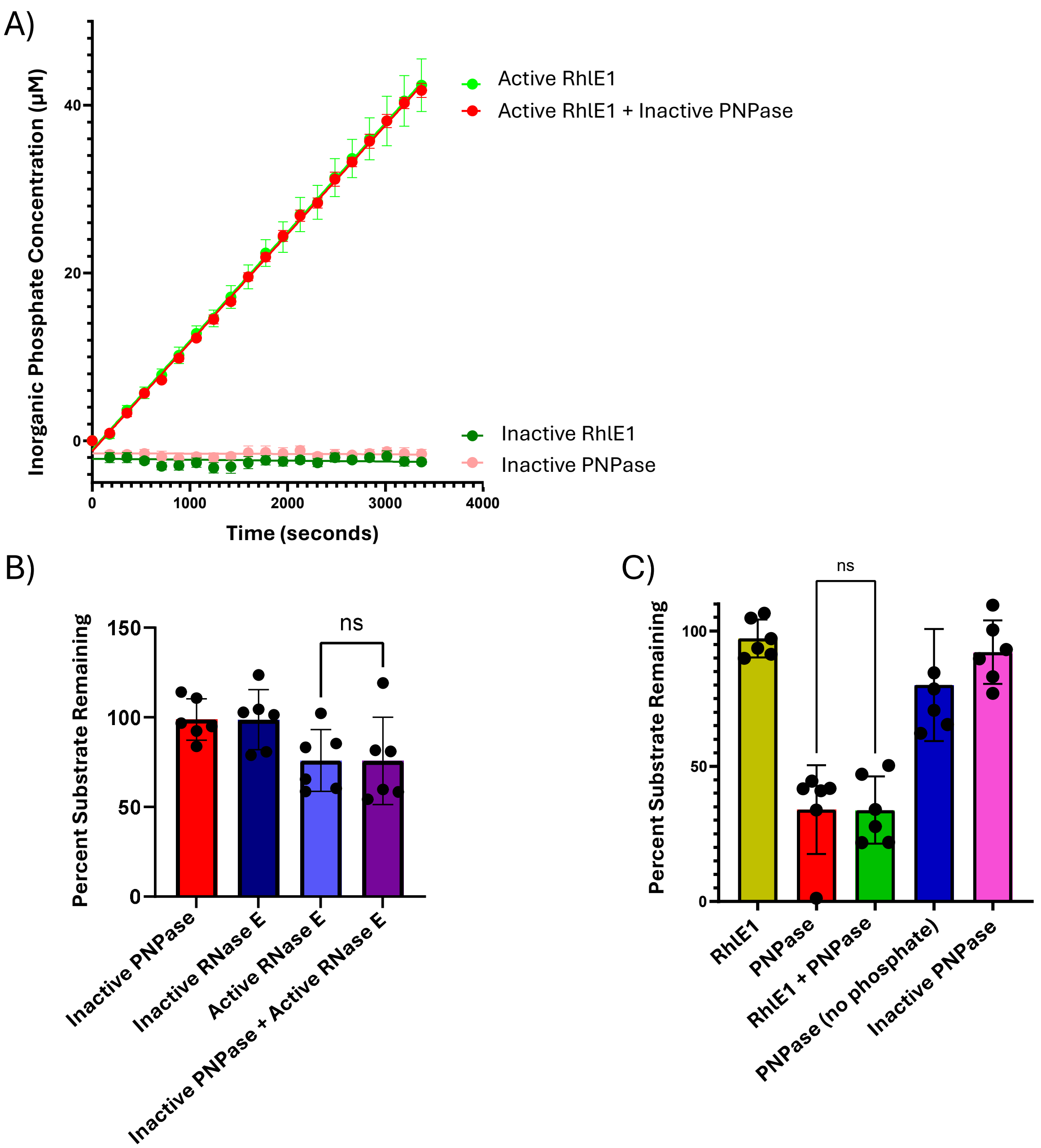


**Supplementary Figure 9.** Testing for allosteric activation of degradosome enzymes. **A)** Phosphate production assay of RhlE1 in combination with catalytically inactive PNPase. Points represent the average of three replicates and simple linear regression was performed. Error bars represent the standard error of the mean for each time point. **B)** RNA cleavage assay quantification of catalytically active full-length RNase E and catalytically inactive PNPase. **C)** RNA cleavage assay quantification of catalytically active PNPase and RhlE1. For cleavage assays, the band intensities of six replicates were quantified on ImageJ with the FIJI plugin and shown with dots, and the average is shown with the bar. Error bars represent the standard error of the mean. An unpaired t-test was performed for significance testing. ns $p > 0.05$

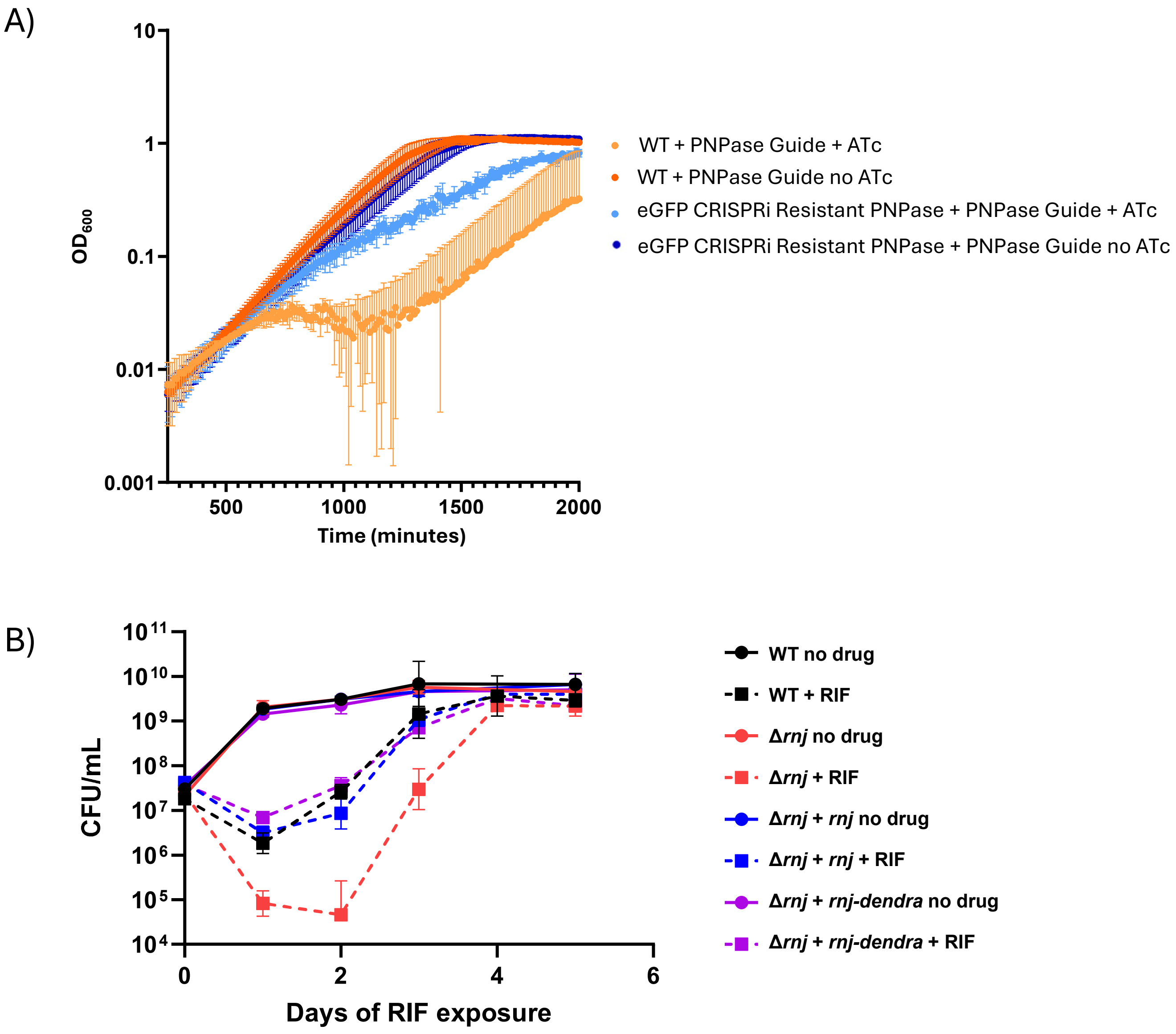


**Supplementary Figure 10. A)** Growth curve of PNPase tagged with eGFP. CRISPRi knockdown of the native PNPase in the +ATc (50 ng/mL) condition. **B)** Kill curve with rifampicin (12 µg/mL) to test the functionality of RNase J-dendra2 .

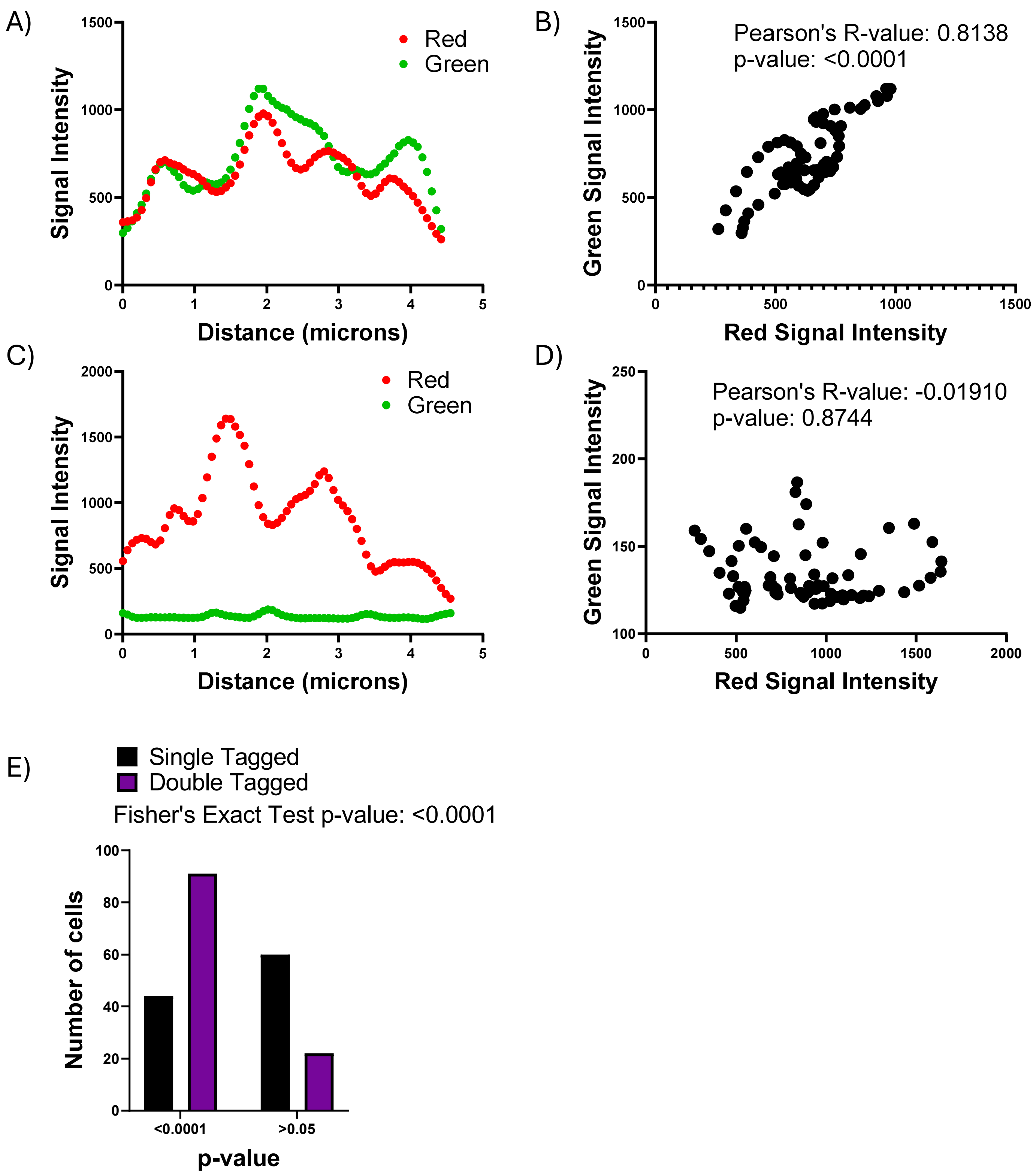


**Supplementary Figure 11.** Examples to demonstrate microscopy quantification methods. A line was drawn through the long axis of each cell in FIJI, and the signal intensity was obtained. For double-tagged fluorescent strains, localization was assessed using Pearson's R-value. A plot of the intensity versus distance is shown in **A)** and a plot of the green signal intensity versus red signal intensity is shown in **B)** for a cell in which the signals were localized. **C)** and **D)** show the same types of graphs as B) and C), respectively, for a cell in which the signals were not localized. The Pearson's R-values are shown in this figure and were reported in Figures 6A-F and Figure 7B-G. **E)** shows the p-values obtained from Pearson's R-value calculations as in B) and D) for 150 cells. p-values were reported if they were below 0.0001 or above 0.05, and a Fisher's Exact test was used to assess the statistical difference between the single-tagged and double-tagged groups.

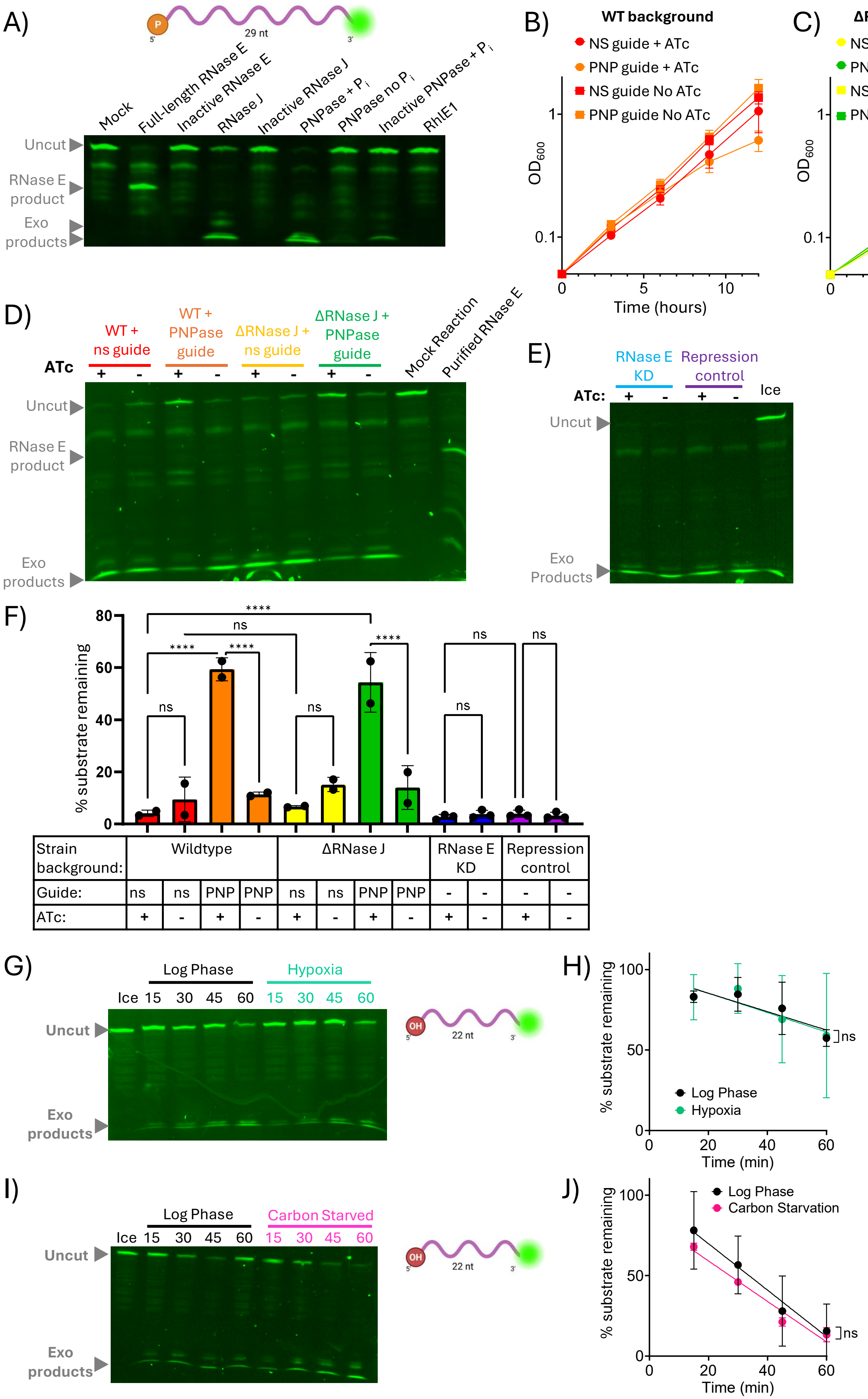


| Strain background: | Wildtype | | | | ΔRNase J | | | | RNase E KD | | Repression control | |
|---|---|---|---|---|---|---|---|---|---|---|---|---|
| Guide: | ns | ns | PNP | PNP | ns | ns | PNP | PNP | - | - | - | - |
| ATc: | + | - | + | - | + | - | + | - | + | - | + | - |

**Supplementary Figure 12.** **A)** Cleavage assay of purified *M. smegmatis* degradosome enzymes and a 29 nt fluorescently labeled RNA. **B-C)** Growth curves using the CRISPRi system to knockdown PNPase (PNP) or a nonspecific (NS) guide control in B) the wildtype background or C) the RNase J knockout background. **D-E)** *M. smegmatis* lysates were incubated with the 29 nt labeled RNA used in A). WT and RNase J deletion strains with and without CRISPRi knockdown of PNPase for 9 hr are shown in D), with cleavage of the same oligo by purified RNase E shown as a control. A tet-repressible RNase E knockdown strain described in Zhou et al 2024 and its matched control strain were used in E). “Ice” indicates incubation of RNA with a WT or control strain lysate on ice. **F)** Quantification of replicate assays as shown in D-E). Ordinary one-way ANOVA was used for significance testing with Sidak’s multiple comparisons test. **G-J)** A 22 nt fluorescently labeled RNA was incubated with lysates from cultures growing in log phase or exposed to the indicated stress conditions. The oligo is expected to be cleaved by RNase J and PNPase but not by RNase E (Rapiejko et al 2026). Simple linear regression of the lines of best fit were compared for significance testing. Points represent the average of two or three biological replicate strains. ns $p > 0.05$, **** $p \leq 0.0001$
